# Latrophilin-2 mediates flow activation of Notch to control vascular endothelial phenotype and atherosclerosis

**DOI:** 10.1101/2025.07.01.662676

**Authors:** Keiichiro Tanaka, Minghao Chen, Divyesh Joshi, Manasa Chanduri, Yoo-Min Koh, Xiaoli Wang, Hanming Zhang, Xing Zhang, Jay Solanki, Carlos Fernandez-Hernando, Martin A. Schwartz

## Abstract

Fluid shear stress (FSS) is a major determinant of endothelial cell (EC) phenotype, with physiological laminar FSS promoting arterial identity and stability, and disturbed FSS increasing susceptibility to atherosclerosis. We previously identified the adhesion G protein-coupled receptor (GPCR) Latrophilin-2 (Lphn2) as a junctional protein required for FSS activation of the PECAM1/VE-cadherin/VEGFR2/PlexinD1 junctional mechanosensory pathway, which regulates EC alignment to laminar blood flow and promotes EC inflammatory activation and atherogenesis under disturbed flow. Here, we demonstrate that *Lphn2* endothelial cell-specific knockout hyper-lipidemic mice develop larger atherosclerotic plaques than controls—opposite from the previously reported *Pecam1* knockout model. Transcriptomic analysis revealed that Lphn2 contributes nearly half of flow-responsive gene expression and suppresses pro-inflammatory pathways. Critically, Lphn2 is essential for FSS-induced Notch1 activation to suppress inflammation, via a GPCR-in-dependent mechanism. Active Notch1 correlates with physical association of Lphn2 with Notch1 and its activator, γ-secretase, in response to flow. Both the Notch1 intracellular domain (N1ICD) and transmembrane domain (N1TMD) contribute to the anti-inflammatory gene program, with the non-canonical N1TMD signaling specifically suppressing the pro-inflammatory YAP pathway. Together with the accompanying paper showing that Lphn2 also mediates flow activation of the Smad1/5 pathway, these data identify Lphn2 as a central mediator of EC flow responses through multiple independent mechanisms in vascular biology and disease.

## Introduction

Atherosclerosis, a chronic inflammatory disease of arterial walls, is the leading cause of morbidity and mortality worldwide^1,2^. Lesions form preferentially at arterial branch points and regions of curvature where endothelial cells (ECs) are exposed to low and multidirectional flow patterns, termed disturbed FSS, whereas regions of high laminar shear stress are protected^3,4^. Flow-sensing mechanisms are thus crucial in vascular health and disease.

Latrophilin-2 (Lphn2) is a member of the adhesion GPCR family that is well expressed in vascular endothelial cells^5^. Adhesion GPCRs have large extracellular domains that bind ligands on other cells or extracellular matrices and contains a G protein autoproteolysis (GAIN) domain that generates an internal peptide that mediates G protein activation^6–8^. We recently reported that Lphn2 is required for activation of the PECAM-1/VE-Cadherin/VEGFR/Plexin D1 junctional mechanosensory complex by FSS^5,9,10^. Lphn2 appears to respond to FSS-induced changes in plasma membrane composition or fluidity^5,11^. Lphn2 was found to be essential for activation of early flow-induced signals, for EC alignment in flow and for flow-dependent vascular remodeling in mice and zebrafish^5^. In addition, Lphn2 has been implicated in maintaining endothelial barrier integrity and regulating vascular permeability^5,12^, as well as angiogenesis^5,13^ and the response to oxidative stress^14^. However, its roles in vascular inflammation and atherosclerosis were not determined.

EC Notch signaling regulates vascular development, homeostasis, and disease^15^. Notch1 is the main Notch receptor in ECs and is activated by binding ligands such as DLL4 and Jag1 on adjacent cells^16^, which triggers Notch activation through proteolysis by ADAM10 and gamma-secretase. FSS amplifies ligand-dependent Notch activation through unknown mechanisms^17,18^. The released Notch1 intracellular domain (N1ICD) translocates to the nucleus, binds the transcription factor RBPJ and drives expression of target genes, including HES family genes, such as Hes1^19^. This canonical Notch signaling pathway inhibits VEGF receptor expression and angiogenesis, induces quiescence and promotes arterial identity^20–22^. Notch1 cleavage also releases its transmembrane domain (N1TMD), which remains in the plasma membrane where it associates with VE-cadherin to stabilize EC cell-cell junctions and improve barrier function (non-canonical signaling)^23^. In contrast, atherogenic factors, including inflammatory cytokines, dyslipidemia and disturbed shear stress, inhibit Notch signaling^24–26^. Endothelial specific knockout (ECKO) of Notch1 exacerbates atherosclerosis, while its activation is protective^25^, consistent with an anti-inflammatory, stabilizing function.

We began this study by investigating the role of Lphn2 in atherosclerosis. Based on its position upstream of the junctional mechanosensory complex that promotes EC inflammatory activation in disturbed flow^27^, Lphn2 was expected to facilitate inflammatory gene expression. However, Lphn2 ECKO increased atherosclerotic plaque size and suppressed pro-inflammatory genes. Prompted by their co-localization at junctions and Notch’s anti-inflammatory effects, we investigated possible connections between Lphn2 and Notch1. These studies showed that Lphn2 is required for activation of Notch by FSS and elucidated protein physical associations and functional consequences of the Lphn2-Notch axis.

## Results

### Latrophilin-2 endothelial-specific knock-out increases atherosclerotic plaques in mice

Previous studies showed that deletion of PECAM-1 in mice, a component of the junctional mechanosensory complex, reduced inflammation and atherosclerosis at regions of disturbed FSS^27–29^. To investigate the contribution of Lphn2 to atherosclerosis, we generated EC-specific *Lphn2* knockout mice (*Lphn2* ECKO) using a tamoxifen-inducible Cre-loxP system, with Cre recombinase driven by the Cdh5 (VE-cadherin) promoter. Knockdown was confirmed as shown in Supplemental Figure 1A. To induce hyperlipidemia, mice were injected with adeno-associated virus (AAV) encoding Pcsk9 that reduces hepatic LDL receptor protein, then fed a high-fat diet (HFD; Figure 1A). *Lphn2* ECKO significantly increased atherosclerotic plaque size in both male and female mice (Figure 1B,1C and Supplemental Figure 2A and 2B) without altering circulating cholesterol or triglycerides (Figure 1D). *Lphn2* ECKO mice also showed larger plaques in aortic root sections (Figure 1E and 1F), with slightly greater collagen accumulation, less smooth muscle cell recruitment, and no significant change in CD68+ macrophage content (Supplemental Figure 2C-G). Consistent with these findings, analysis of a public single-cell RNA sequencing dataset from human carotid arteries revealed enrichment of *LPHN2*, but not *LPHN1* and *LPHN3*, in plaque endothelial cells (Supplemental Figure 3). Lphn2 in ECs thus restrains atherosclerosis, opposite from PECAM-1.

**Figure 1:**
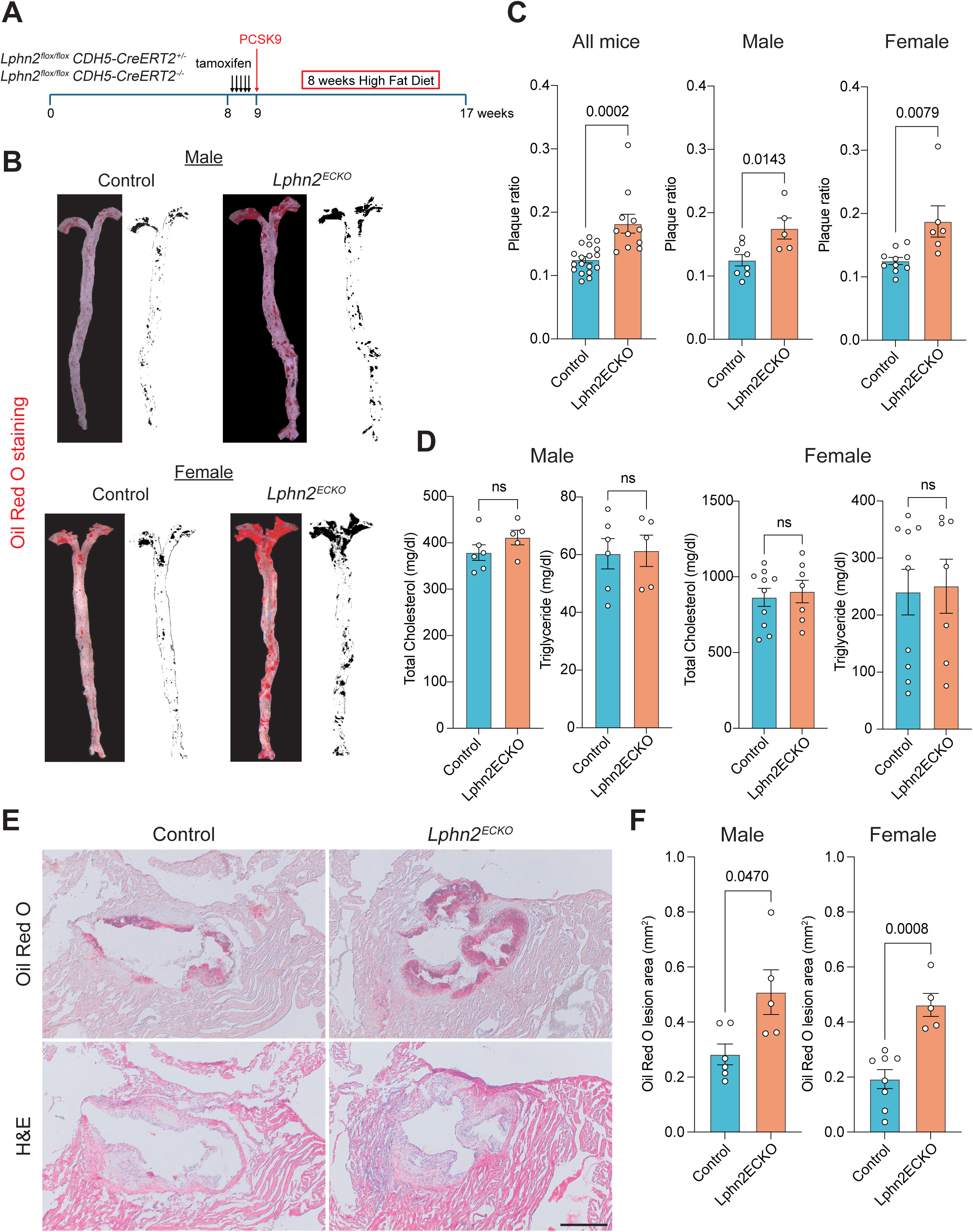
Lphn2 endothelial-specific knockout increases atherosclerosis. (**A**) Experimental timeline: Control or *Lphn2* ECKO 8-week-old mice were treated with tamoxifen and subsequently injected with PCSK9 AAV8 virus. Mice were fed a high-fat diet for 8 weeks, then tissue was harvested. (**B**) Representative images of Oil Red O-stained aortas and corresponding plaque masks from control and *Lphn2* ECKO mice (male and female). (**C**) Quantification of atherosclerotic plaque area in *Lphn2* ECKO mice versus control mice (plaque area ratio to total area). Data were from 7 control males, 5 *Lphn2* ECKO males, 10 control females and 6 Lphn2 ECKO females. Statistics were calculated using unpaired *Student*’s t-test. (**D**) Quantification of blood total cholesterol and triglyceride levels in mice from (**C**). Statistics used unpaired *Student*’s t-test. (**E**) Oil Red O and hematoxylin and eosin (H&E) staining of aortic root sections from control and *Lphn2* ECKO mice. Scale bar: 500µm. (**F**) Quantification of Oil Red O lesion area from control and *Lphn2* ECKO mice. Statistics: unpaired *Student*’s t-test.

### Lphn2 in flow-responsive gene regulation

To further investigate the role of Lphn2 in endothelial cells shear stress responses, we analyzed transcriptomic effects of Lphn2 knockdown in HUVECs under physiological laminar shear stress (12 dynes/cm^2^) (Figure 2A). Lphn2 knockdown significantly altered gene expression under static conditions compared to control siRNA (Figure 2B), but had a much stronger effect under flow, blocking nearly half of the flow-induced changes in gene expression (Figure 2C). Among flow-responsive genes that did not require Lphn2, Klf2 and Klf4 were among the top hits (Figure 2D), consistent with prior findings^5^. For comparison, we knocked down another important mechanosensor protein, PIEZO1 (Supplemental Figure 4A; knockdown confirmation in Supplemental Figure 1B). *PIEZO1* knockdown affected about a third as many flow-responsive genes (Supplemental Figure 4B). Interestingly, genes affected by *PIEZO1* vs *LPHN2* knockdown showed no significant overlap (Supplemental Figure 4C). According to KEGG pathway enrichment analysis, *LPHN2* knockdown increased multiple inflammatory pathways with the TNFα/NF-κB pathway as the top upregulated pathway. The Hippo pathway was the top downregulated pathway (Figure 2E), consistent with Hippo pathway kinases suppressing the Yap and Taz transcription factors that promote EC inflammatory gene expression and atherosclerosis in mice^30–32^. These transcriptional shifts suggest that loss of Lphn2 enhances EC pro-inflammatory phenotypes, consistent with increased atherosclerosis in *Lphn2* ECKO mice.

**Figure 2:**
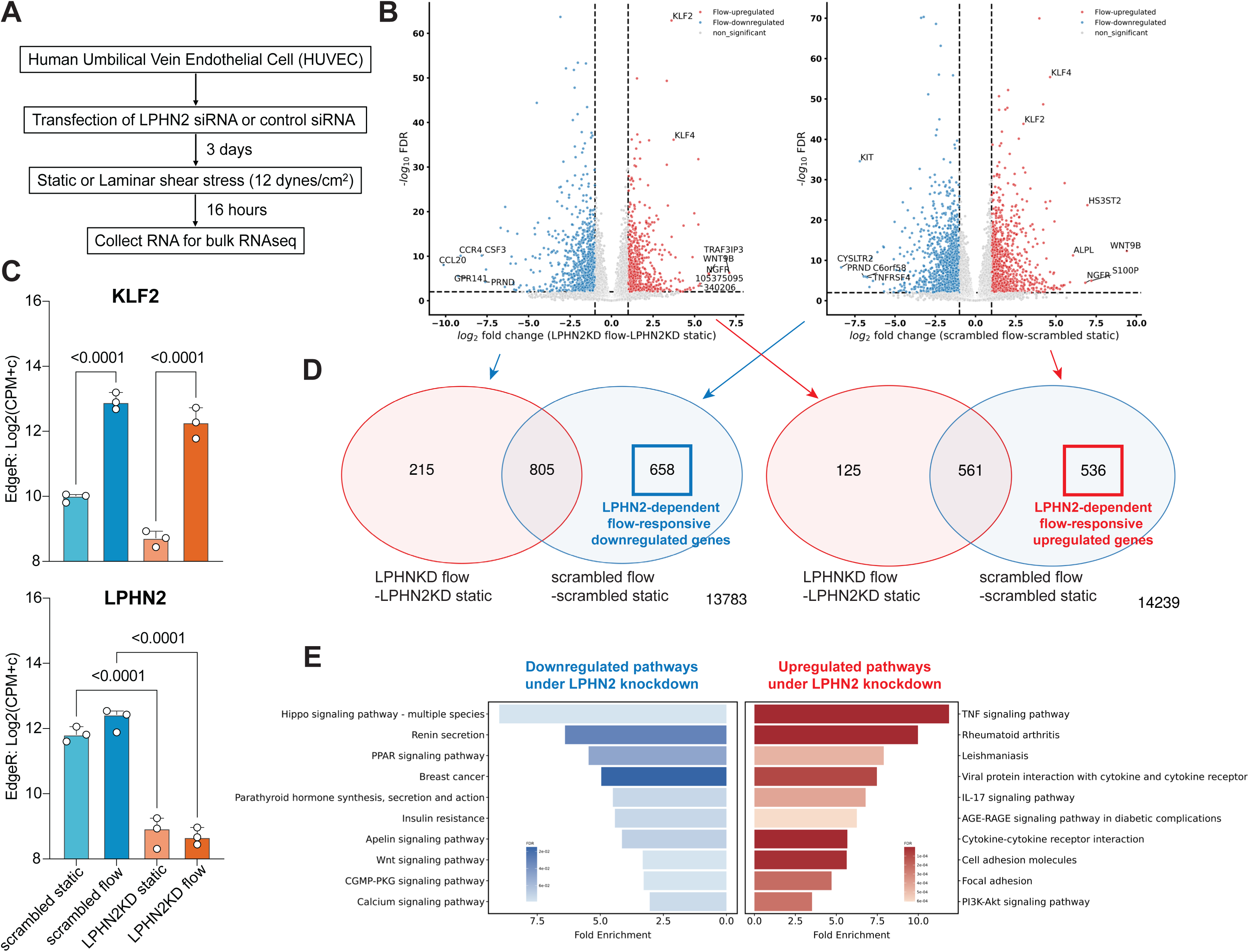
LPHN2-dependent regulation of flow-responsive transcriptome. (**A**) Schematic of experimental procedure. Human umbilical vein endothelial cells (HUVECs) were transfected with control or *LPHN2* siRNAs. After 3 days, cells were subjected to laminar shear stress at 12 dynes/cm^2^ for 18 h or maintained under static (no flow) conditions. Total RNA was extracted and analyzed by bulk RNAseq. (**B**) Volcano plots of log_2_ fold change (flow vs static) versus -log_10_ FDR showing flow-responsive differentially expressed genes (DEGs) in LPHN2 vs control siRNAs. The top 5 upregulated or downregulated DEGs and *KLF2/KLF4* are highlighted. (**C**) Venn diagram illustrating overlaps of flow-responsive downregulated and upregulated DEGs in control and *LPHN2* knockdown cells from (**B**). The number on the lower right indicates the number of genes that responded to flow under control but not under *LPHN2* KD. (**D**) Normalized expression counts for *KLF2* and *LPHN2* analyzed by edgeR. Statistical significance was determined by one-way ANOVA with Tukey post-hoc analysis. (**E**) KEGG pathway enrichment for flow-responsive LPHN2 target genes identified in (**D**). X-axis is fold enrichment and color scale indicates FDR, with pathways upregulated by *LPHN2* KD in red and downregulated pathways in blue.

### Lphn-2 Is required for flow-induced Notch activation

The inconsistency between these results and the established role of the junctional mechanosensory pathway in shear stress signaling and atherosclerosis led us to investigate whether other shear stress pathways might be regulated by Lphn-2. Given that Lphn-2 and the Notch pathway are activated by shear stress at cell-cell junctions, and Notch signaling protects against atherosclerosis^17,25^, we considered whether Lphn2 might regulate Notch. FSS induces both canonical Notch signaling through its intracellular domain that regulates gene expression and non-canonical signaling through its transmembrane domain that stabilizes cell-cell junctions^23^. We therefore silenced *LPHN2* and assessed flow-induced Notch activation. *LPHN2* KD markedly reduced N1ICD production in response to shear stress (Figure 3A–3B and Supplemental Figure 1C). By contrast, its depletion had no effect on Notch activation by its ligand DLL4 (Figure 3C–3D). *LPHN2* depletion also attenuated flow induction of the canonical N1ICD direct target gene Hes1 in both HUVECs and HAECs (Figure 3E–3G). Consistent with *in vitro* findings, *en face* analysis of the thoracic aorta from *Lphn2* ECKO mice revealed reduced Hes1 protein (Figure 3H-3I). Together, these data demonstrate a requirement for Lphn2 in flow-induced Notch pathway activation *in vitro* and *in vivo*.

**Figure 3:**
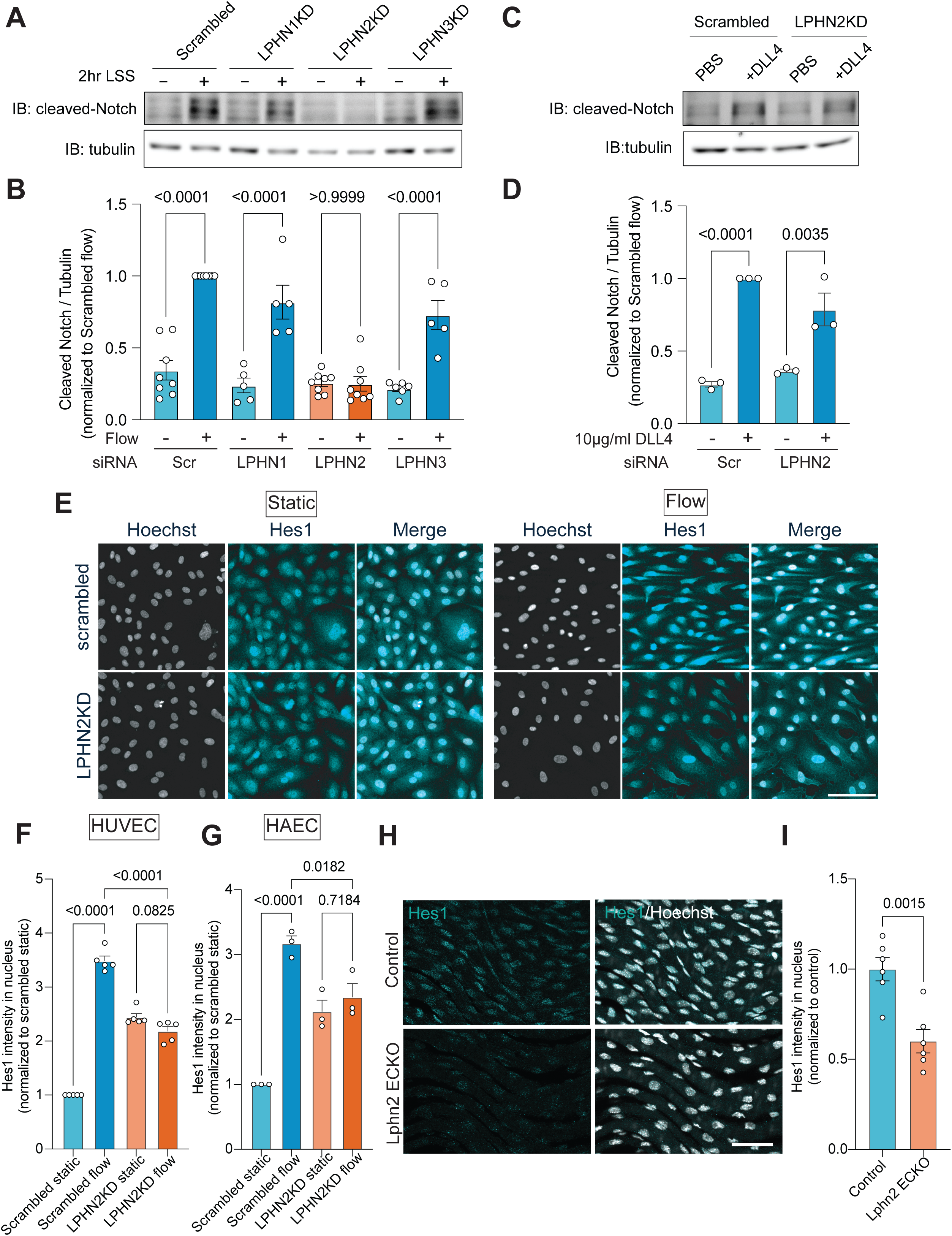
LPHN2 is required for flow-induced Notch1 activation. (**A**) ECs exposed to shear stress, with knockdown (KD) of Lphn 1-3 or control were lysed and assayed by Western blot for cleaved Notch1 intracellular domain (N1ICD) to assess Notch1 activation. (**B)** Quantification of N1ICD in (**A**). Statistics analyzed using one-way ANOVA with Tukey post-hoc analysis. (**C**) Notch activation in response to DLL4 ligand in control and *LPHN2* KD cells. **(D)** Quantification of N1ICD levels from (**C**). Data were from 3 independent experiments. Statistics used one-way ANOVA with Tukey post-hoc analysis. (**E**) Representative images of Hoechst and Hes1 staining of HUVECs with or without laminar shear stress for 16 hours. Scale bar: 50µm. (**F**) Quantification of nuclear Hes1 intensity from (**E**). Quantifications were from 5 independent experiments. Statistics used one-way ANOVA with Tukey post-hoc test. (**G**) Hes1 nuclear intensity in human aortic endothelial cells (HAECs) as in (**F)**. Data were from 3 independent experiments. Statistics used one-way ANOVA with Tukey post-hoc. (**H**) *En face* immunostaining of thoracic aorta from *Lphn2* ECKO and control mice for Hes1. (**I**) Quantification of Hes1 nuclear intensity from (**H**). N = 6 mice per condition. Statistics were analyzed using unpaired *Student*’s t-test.

### Notch activation by flow is independent of G Proteins and beta-arrestins

We next investigated the role of G proteins in LPHN2-dependent activation of Notch (Schematic in Figure 4A). Knockdown of the G proteins involved in endothelial flow signaling had no effect on flow-induced Notch1 cleavage (Figure 4B–4C). To independently test this result, we rescued endogenous *LPHN2* knockdown with wild-type *LPHN2* vs the GPCR-defective mutant (H1071A)^33^. Both constructs restored flow-induced Hes1 expression equally well (Figure 4D–4E), supporting G protein-independent function. We additionally assessed β-arrestins, which are key effectors in G protein-independent GPCR signaling. Knockdown of β-arrestin 1 (ARRB1), the major isoform in ECs, had no effect on flow-induced Hes1 nuclear enrichment (Supplemental Figure 5A-5C). We noticed that *ARRB1* KD elevated baseline Hes1 expression, likely due to the reported impairment of Notch1 degradation^34^. Lphn2-dependent Notch activation thus appears to be β-arrestin-independent.

**Figure 4:**
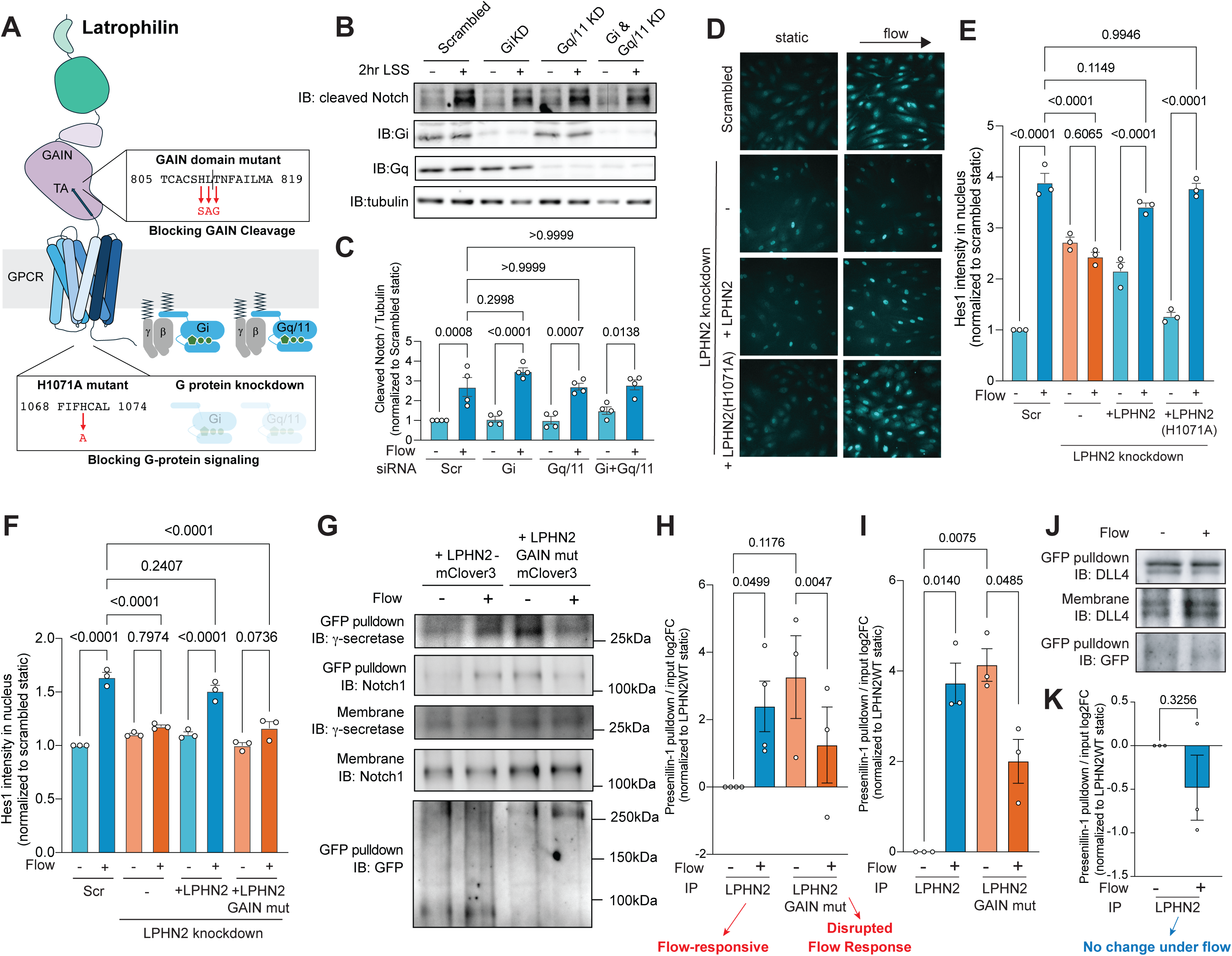
LPHN2 mediated Notch1 activation is G protein-independent but GAIN autoproteolysis-dependent. (**A**) Schematic of Latrophilin-2 (LPHN2) structure with mutations indicated. (**B**) EC lysates Western blotted for flow-induced NICD after knockdowns of the G proteins that couple with LPHN2. (**C**) Quantification of NICD levels from (**B**). Statistical analysis used one-way ANOVA with Tukey post-hoc analysis. (**D**) Representative images of Hes1 staining following rescue with wildtype or GPCR-deficient LPHN2. (**E**) Quantification of Hes1 nuclear intensity from (**D**). Data were from 3 independent experiments. Statistics used one-way ANOVA with Tukey post-hoc analysis. (**F**) Hes1 expression in ECs after rescue of LPHN2 knockdown with WT vs. LPHN2 GAIN domain autoproteolysis mutant. Data were from 3 independent experiments. Statistics used one-way ANOVA with Tukey post-hoc analysis (**G**) Co-Immunoprecipitation of WT LPHN2 or GAIN domain mutant with γ-secretase and Notch1 using isolated membrane fractions as inputs. γ-secretase was detected using an antibody targeting Presenillin-1. (**H&I**) Quantification of co-IPs of LPHN2/ɣ-secretase and LPHN2/Notch1 from (**G**). Values are log2 fold changes compared to LPHN2 wild-type static. Statistics used paired *Student*’s t-test. (**J**) Co-IP of LPHN2 and DLL4 under static vs. flow conditions. (**K**) Quantification from (**J**) as in (**H&I**). Statistics used paired *Student*’s t-test.

**Figure 5:**
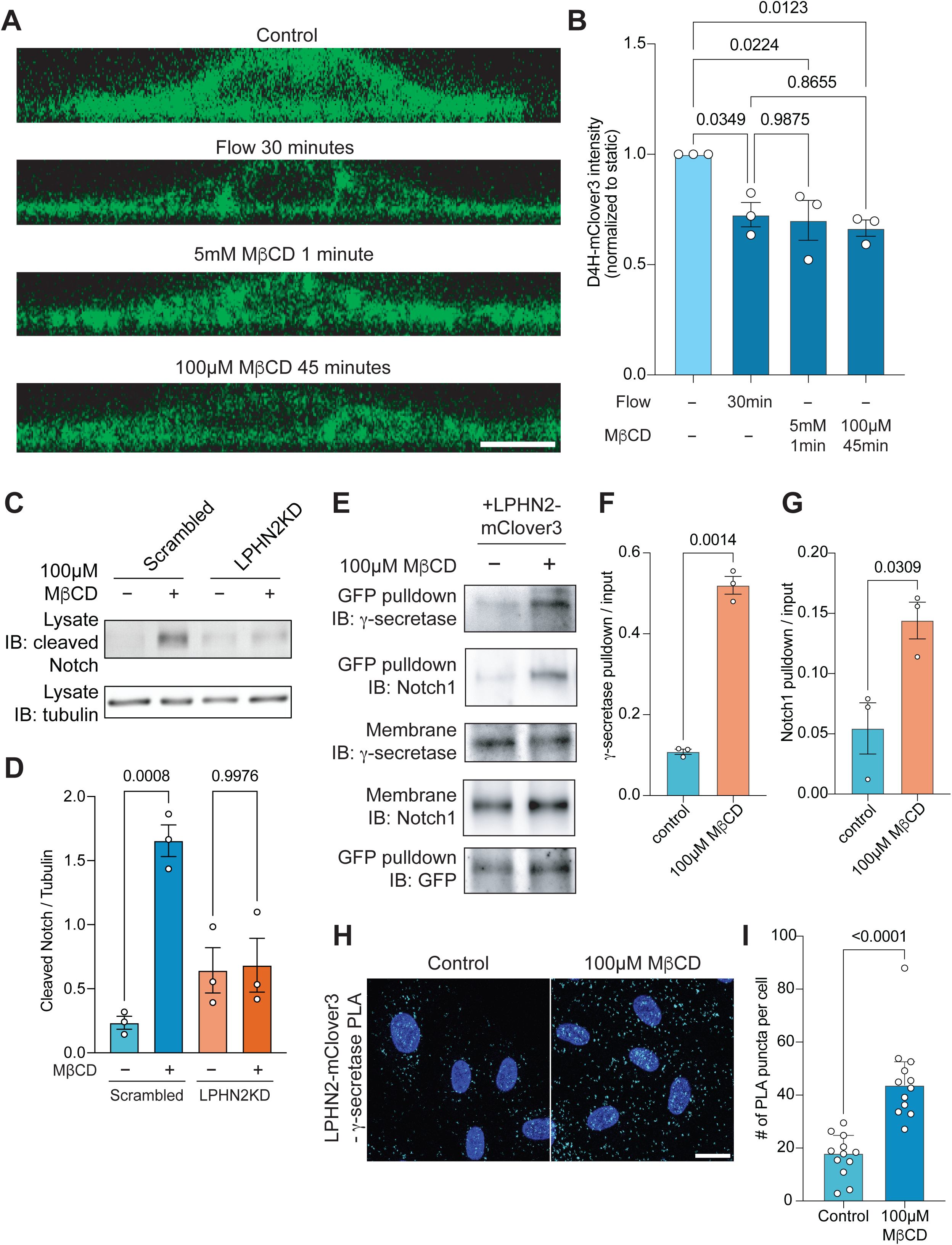
LPHN2 mediates membrane cholesterol-dependent Notch1 activation. (**A**) Representative XZ projections of the D4H-mClover3 cholesterol reporter in HUVECs under the indicated conditions. Scale bar: 10µm. (**B**) Quantification of D4H-mClover3 fluorescence intensity from cell membrane regions under the indicated conditions. Data were from 3 independent quantifications. Statistics used one-way ANOVA with post-hoc Tukey. (**C**) Immunoblot for N1ICD in control vs *LPHN2* KD ECs ± MβCD treatment . (**D**) Quantification of N1ICD levels from (**C**). Statistics: one-way ANOVA with Tukey post-hoc test. (**E**) ECs expressing Lphn2-mClover3 were treated ± MβCD, lysed, mClover3 immunoprecipitated and Westerns probed for Notch1 and ɣ-secretase component, Presenillin-1. (**F&G**) Quantification of co-IPs from (**E**). Statistics used unpaired *Student*’s t-test. (**H**) Representative images of proximity ligation assay (PLA) targeting LPHN2-mClover3 and γ-secretase component, Presenillin-1. DAPI staining and PLA puncta are shown. Scale bar: 20µm. (**I**) Quantification of PLA puncta from (H). Statistics calculated using unpaired *Student*’s t-test.

### LPHN2 GAIN domain mutant and physical association with Notch components

To further investigate the mechanism of Lphn2-dependent Notch signaling, we examined an *Lphn2* mutant lacking GAIN domain autoproteolytic activity (see Figure 4A). This mutant failed to support flow-induced Notch pathway activation (Figure 4F). We next investigated whether LPHN2 physically interacts with any components of the Notch pathway. For these experiments, we expressed mClover3-tagged wild-type (WT) or GAIN domain mutant LPHN2 and examined cells with or without FSS for 2 hours. Immunoprecipitation of isolated membrane fractions using GFP-TRAP beads followed by Western blotting showed a flow-induced increase in the association of WT LPHN2 with Notch1 and γ-secretase, the enzyme responsible for the terminal Notch1 cleavage that generates the N1ICD and N1TMD. By contrast, the GAIN mutant showed the opposite behavior, binding was high without flow and decreased with FSS (Figure 4G–4I). As a control, we assayed the association of Notch1 with its ligand DLL4. These proteins co-immunoprecipitated but displayed no change after flow (Figure 4J-4K). Lphn2-dependent Notch activation by flow thus requires an intact GAIN domain, which correlates with flow-induced physical associations.

### LPHN2 responds to changes in membrane fluidity to activate Notch

Shear stress has been reported to rapidly increase plasma membrane fluidity associated with decreased cholesterol content^11,35^. We previously reported that Lphn2 is required for EC activation of the PECAM1 pathway in response to increasing plasma membrane fluidity, induced by depleting membrane cholesterol with methyl-β-cyclodextrin (MβCD)^5^. To test whether membrane fluidity mediates flow-induced Notch activation, we treated ECs with a low concentration of MβCD to reduce membrane cholesterol by an amount similar to FSS, demonstrated using the D4H* cholesterol reporter (Figure 5A–5B). Removing cholesterol with MβCD activated the Notch pathway similarly to shear stress, which was blocked by *LPHN2* depletion (Figure 5C–D). Removing cholesterol also enhanced the co-immunoprecipitation of LPHN2 with γ-secretase, which was detected by Presenillin-1, and Notch1 (Figure 5E-G). Consistent with this, proximity ligation assays demonstrated a significant increase in binding between LPHN2 and γ-secretase following cholesterol removal by MβCD (Figure 5H-I). We also tested the effect of cholesterol extraction on induction of the Notch1 target gene *HES1*. Initial experiments showed that transferring ECs to the very low serum medium required for effective cholesterol extraction by MβCD was sufficient to increase baseline HES1 expression; we therefore used HPβCD (2-Hydroxypropyl-β-cyclodextrin), which can be used in serum-containing medium^36^. Cholesterol extraction in this condition induced HES1 expression, which was abrogated by *LPHN2* knockdown, supporting Lphn2-dependent Notch1 activation (Supplemental Figure 6). Together, these data show that changes in membrane fluidity drive shear stress activation of Notch through LPHN2, likely through changes in intramembrane interactions.

### Role of Notch1 in *LPHN2*-dependent gene expression

To further assess the role of Notch signaling downstream of *LPHN2*, we evaluated the role of the canonical Notch1 intracellular domain (N1ICD) signaling vs noncanonical transmembrane domain (N1TMD). To address this question, we rescued Lphn2 knockdown by overexpressing either the N1ICD or the N1TMD (Figure 6A-6C and Supplemental Figure 7A). N1ICD overexpression upregulated multiple Notch target genes, including Hes5 by 3.3-fold, HeyL by 9.7-fold, and Hey2 by 2.1-fold (Supplemental Figure 7B). Quantification of differentially expressed genes (DEGs) revealed that approximately 30% the Lphn2-dependent genes were reversed by expression of the N1TMD, while 22% were reversed by expression of the N1ICD (Figure 6C and 6E and Supplemental Figure 7C-7E). For genes rescued by N1TMD, KEGG pathway enrichment analysis identified the Hippo pathway as the most downregulated by LPHN2 KD, while multiple inflammatory pathways were upregulated (Figure 6D). As Hippo suppresses inflammatory Yap/Taz signaling, these results all point toward an anti-inflammatory role for the N1TMD. For N1ICD over-expression, the PPAR signaling and categories related to metabolism, cell cycle and inflammation were most affected (Figure 6F). Given the effects of LPHN2 on inflammatory gene expression and canonical YAP targets (Figure 2 and Supplemental Figure 7F) and the paucity of information about the N1TMD, we further investigated the role of the N1TMD on the Hippo pathway in EC inflammation.

**Figure 6:**
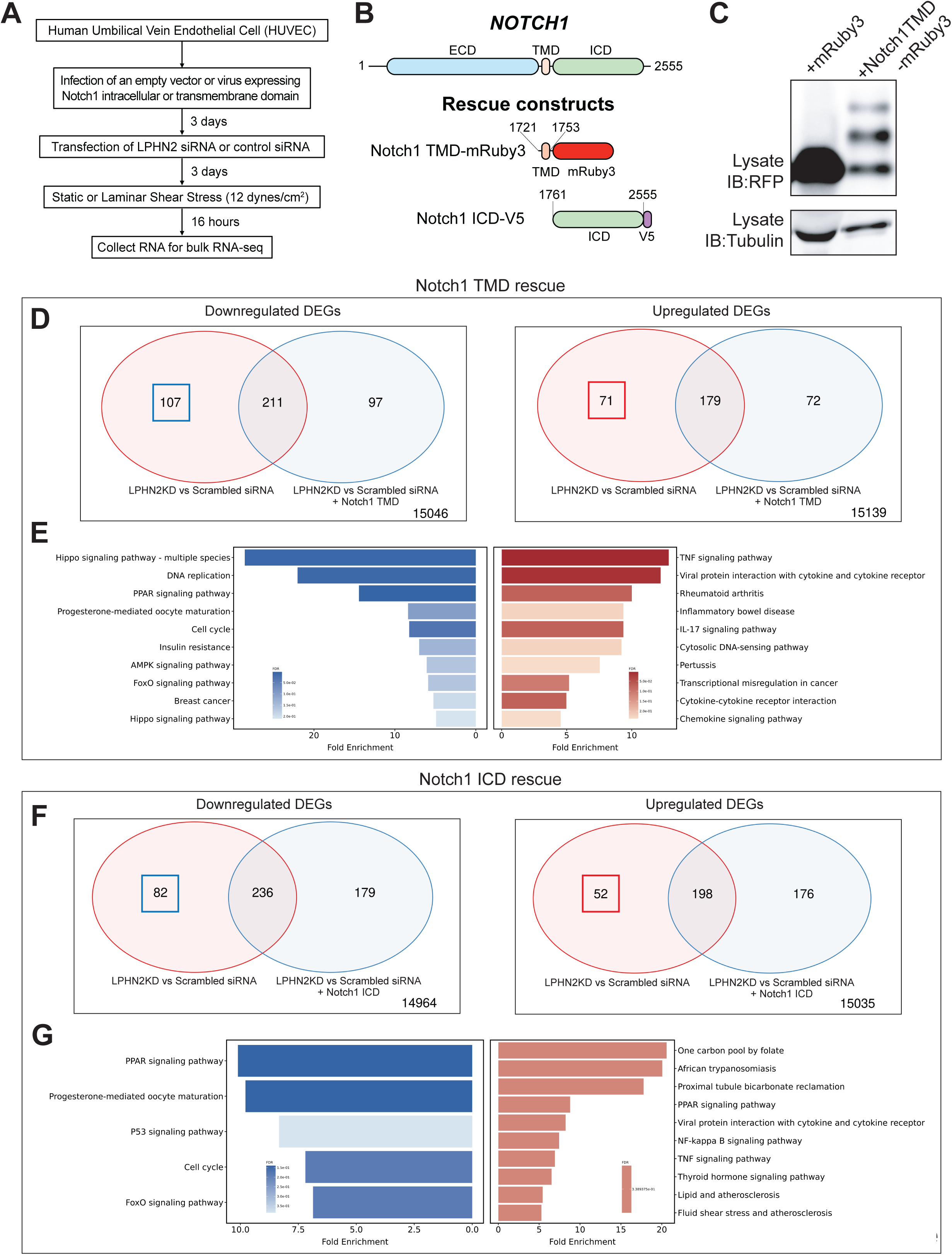
Rescue of LPHN2 knockdowns by Notch1 cleavage products. (**A**) Schematic of experimental procedure. Control and LPHN2 KD HUVECs were infected with lentiviruses encoding Notch1 TMD, Notch1 ICD or empty control, as in Figure 2. (**B**) Schematic of Notch1 rescue constructs. (**C**) Western blot for Notch1 TMD, with tubulin loading control. (**D**) Venn diagrams showing overlap between LPHN2-dependent genes and those rescued by overexpression of Notch1 transmembrane domain (TMD; corresponding volcano plots are shown in Supplemental Figure 6, C and D. (**E**) KEGG pathway enrichment of genes rescued by Notch1 TMD, displayed as in Figure 2E. (**F**) Venn diagram of LPHN2-dependent genes and those rescued by overexpression of Notch1 intracellular domain (ICD; corresponding volcano plots are shown in Supplemental Figure 6, C and E. (**G**) KEGG pathway enrichment analysis of genes rescued by Notch1 ICD.

### Lphn2-Notch Axis Regulates Endothelial Inflammation via YAP transcription factor

The Notch1 TMD was reported to interact with the VE-cadherin TMD to stabilize endothelial junctions and enhance barrier function^23^. As cell-cell adhesions are reported to control Hippo pathway activation^37,38^, we examined effects of the N1TMD on YAP. *LPHN2* knockdown in-creased YAP nuclear localization, while N1TMD overexpression decreased nuclear Yap with or without LPHN2 KD (Figure 7A and 7B). Consistent with the known anti-inflammatory function of Notch signaling^25^, *LPHN2* KD increased basal expression of VCAM1, a marker of endothelial inflammation, which was reversed by the N1TMD (Figure 7C and 7D). A pharmacological inhibitor (K-975) that blocks the interaction of YAP with its downstream transcription factor TEAD also reduced VCAM1 expression, confirming an inflammatory role for YAP following Lphn2 suppression (Figure 7E and 7F).

**Figure 7:**
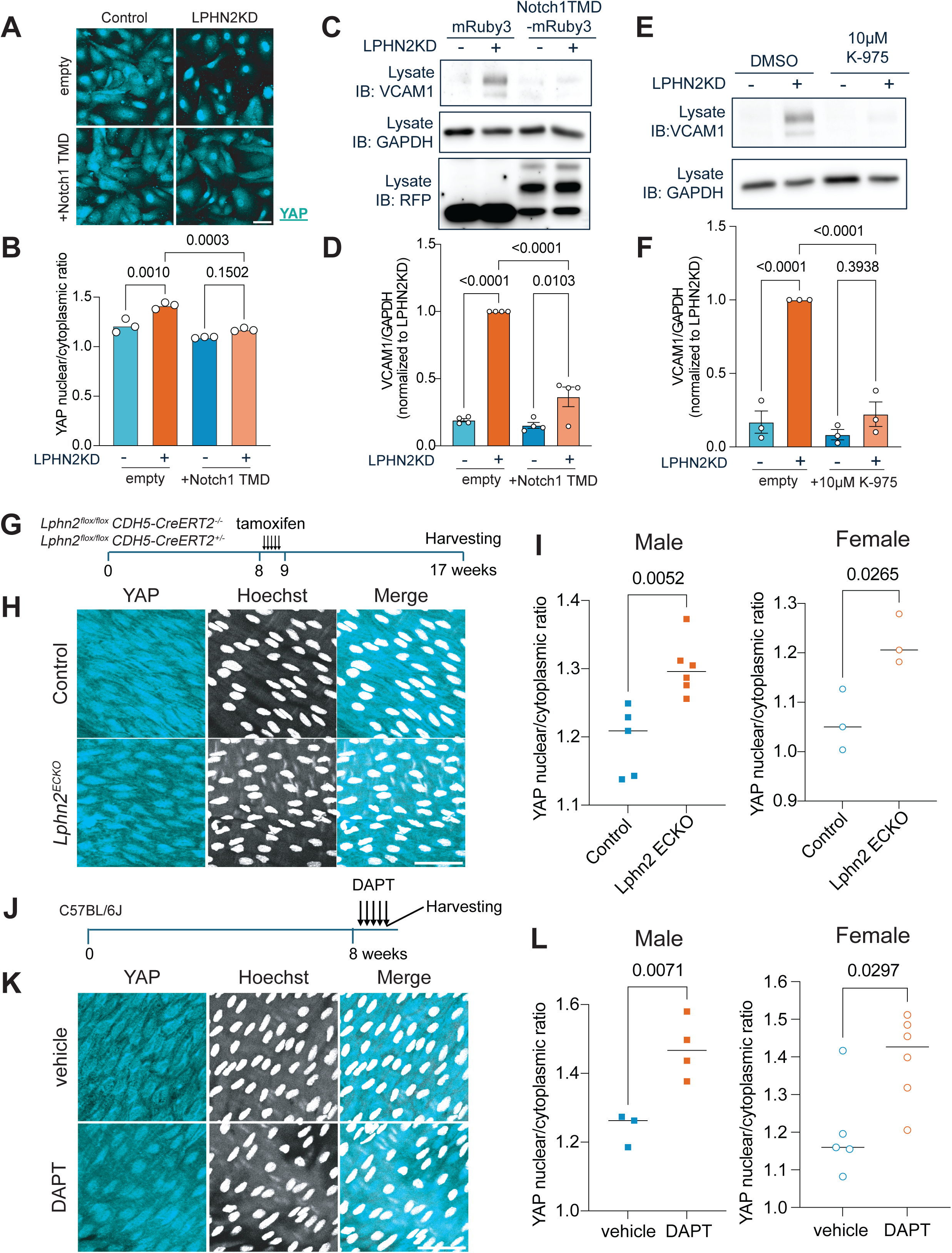
Notch1 TMD regulates YAP. (**A**) YAP nuclear/cytoplasmic ratio in control and *LPHN2* KD cells with or without N1TMD over-expression. (**B**) Quantification of the YAP nuclear:cytoplasmic ratio from (**A**). Data are from three independent experiments. Statistical analysis used one-way ANOVA with Tukey post-hoc test. (**C**) Western blot for VCAM1, an endothelial inflammation marker, in HUVECs with control vs *LPHN2* KD, with or without N1TMD overexpression. GAPDH was used as a loading control (**D**) Quantification of VCAM-1 expression from (**C**). Data are from four independent experiments. Statistical analysis used one-way ANOVA with Tukey post-hoc analysis. (**E**) Western blot for VCAM-1 in control vs *LPHN2* KD cells ± 10 µM YAP-TEAD inhibitor K-975. (**F**) Quantification of VCAM-1 expression normalized to GAPDH from (**E**). Data are from three independent experiments. Statistical analysis used one-way ANOVA with Tukey post-hoc analysis. (**G**) Experimental timeline: Control or *Lphn2* ECKO mice were injected with tamoxifen at 8 weeks of age and tissue harvested 8 weeks post-injection. (**H**) *En face* staining of YAP and Hoechst in control mouse or *Lphn2* ECKO mouse aortae. Scale bar: 50µm. (**I**) Quantification of the YAP nuclear cytoplasmic ratio in endothelial cells from (**H**). Statistics used unpaired *Student*’s t-test. n = 5 (male control), 6 (male *Lphn2* ECKO), 3 (female control), and 3 (female *Lphn2* ECKO). (**J**) DAPT treatment timeline. Control mice over 8 weeks old were injected with 50 mg/kg DAPT or vehicle for 5 consecutive days and tissue was harvested. (**K**) *En face* staining of YAP and Hoechst following DAPT injections or vehicle control. (**L**) Quantification of YAP nuclear/cytoplasmic ratio from (**I**). Statistics used *Student*’s t-test. n = 3 (male vehicle), 4 (male DAPT), 5 (female vehicle), and 6 (female DAPT).

Lastly, we investigated the Lphn2-Notch-YAP axis *in vivo*. *Lphn2* ECKO mice, prepared by injecting tamoxifen into Lphn2^flox/flox^; Cdh5-CreERT2 mice (Figure 7G) showed significantly in-creased nuclear YAP in aortic ECs (Figure 7H and 7I). Pharmacological inhibition of Notch1 cleavage with DAPT also strongly induced YAP nuclear translocation *in vivo* (Figure 7J-L) and *in vitro* (Supplemental Figure 8A and 8B). Collectively, these results demonstrate that pro-inflammatory phenotypes observed in *Lphn2* ECKO mice involve diminished Notch activation, leading to YAP activation.

## Discussion

These results expand the role for Latrophilin-2 in endothelial flow responses and vascular disease beyond regulation of the classical junctional mechanotransduction complex. We first observed larger atherosclerotic plaques in hyperlipidemic *Lphn2* ECKO mice, in contrast to *Pecam-1* KO where plaque size was reduced^27–29^. Transcriptomic analyses revealed that knockdown of Lphn2 blocked approximately 50% of the flow-dependent changes, identifying Lphn2 as a major mediator of flow signaling. In comparison, Piezo1 controls about 1/3 as many genes. Lphn2 knockdown mainly shifted gene expression toward a pro-inflammatory signature, with upregulation of TNFα (NF-κB) and Yap/Taz pathways, which is inconsistent with blockade of the PECAM-1 junctional pathway^9^.

These findings led us to consider the Notch1 pathway, which is activated by FSS, requires cell-cell contact, and promotes EC quiescence and arterial identity^15,17,18^. We found that Lphn2 is required for flow activation of Notch1, as indicated by levels of N1ICD and expression of the target gene HES1. Some of these experiments showed *LPHN2* knockdown slightly increased levels of Hes1 at baseline (i.e., without flow). We did not further investigate this effect but possible explanations include disrupted junctions due to loss of LPHN2, which may indirectly affect Notch activation^5^. The Notch pathway also encodes complex feedback loops that may alter baseline activity^39^.

Flow-induced Notch activation by LPHN2 correlated with a physical association of LPHN2 with both Notch1 and its activator, γ-secretase. Notch appears to control EC function through both its canonical regulation of gene expression by the N1ICD, and non-canonical N1TMD interaction with junctional components. To our surprise, expressed N1TMD had the stronger effect on EC’s transcriptomic profile, reversing ∼30% of the Lphn2-dependent genes, whereas the N1ICD reversed ∼22%. Analysis of DEGs identified the Hippo pathway as the strongest GO category. Hippo pathway kinases inhibit Yap/Taz activation, which promote inflammatory gene expression in ECs and atherogenesis in mouse models. Thus, preventing Notch cleavage should increase nuclear Yap/Taz and upregulate its inflammatory target genes. Indeed, overexpression of the N1TMD suppressed nuclear localization of Yap and expression of the inflammatory marker VCAM-1. These effects are likely due to interaction of the N1TMD with the transmembrane domain of VE-cadherin to stabilize EC junctions^23^, which can control activation of Hippo pathway kinases^31,38^. Thus, the Notch1 TMD likely influences gene expression through effects of junctional organization on Hippo pathway kinases (Supplemental Figure 9).

Lphn2-dependent activation of Notch was independent of G proteins and β-arrestin. However, mutations of the Lphn2 GAIN domain that block autoproteolysis abrogated flow-dependent Notch activation. This result may indicate that the GAIN domain mediates a critical interaction, perhaps even with Notch receptors. However, molecular dynamics simulation of the non-cleavable GAIN domain from ADGRB2 predicted a rigid conformation, while the cleaved one in ADGRL1 is more flexible^40^. Furthermore, in CryoEM structures of Latrophilin-3 holoreceptor, the Latrophilin GAIN domain interacts closely with the 7-membrane spanning TMD^41^. Thus, altering the conformation of the GAIN domain may influence TMD conformations or vice versa. While the GAIN domain could mediate interactions directly, acting indirectly via effects on the transmembrane domain seems equally likely. Indeed, this mechanism fits better with changes in membrane properties as the initiating stimulus. Dissecting these interactions and functions will require a concerted structure-function analysis in future research.

The accompanying paper reports that Lphn2 is also required for flow activation of the Smad1/5 pathway, which is separable from both PECAM1 and Notch pathway activation. In keeping with the major role in flow-dependent gene expression, these results demonstrate that Latrophilin is a central regulator of EC FSS signaling through three independent pathways: the PECAM-1 junctional complex, the Alk1/Endoglin-Smad1/5 pathway, and the Notch pathway, the latter acting through both canonical and non-canonical effectors. The outcome of *Lphn2* ECKO in development, physiology and disease must therefore be due to the sum of the effects on these pathways. In some cases, these effects are opposing so that the Lphn2 phenotype may be milder than otherwise expected. The interactions and effects of Lphns on multiple pathways likely have implications for other cellular or organ systems, especially the central nervous system where both Lphns and Notch play important roles^42,43^. The unique, GPCR-independent mechanisms of Lphn2 signaling in flow may thus offer opportunities to treat disease of the cardiovascular system and elsewhere.

## Methods

### Antibodies

Anti-VCAM1 (abcam) (ab134047); Anti-phospho-p65 (Cell Signaling) (3033); Anti-RFP (Invitrogen) (R10367); Anti-Tubulin (Cell Signaling) (3873); Anti-GAPDH (Cell Signaling) (2118); Anti-YAP (Cell Signaling) (14074); Anti-cleaved Notch (Cell Signaling) (4147); Anti-Hes1 (Cell Signaling) (11988); Anti-Presenillin-1 (Millipore-Sigma) (MAB1563); Anti-Gi (NewEast Biosciences)(26003); Anti-Gq/11 (BD Biosciences) (612705); Anti-Notch1 (Cell Signaling) (4380); Anti-GFP (Invitrogen) (A11122); Anti-DLL4 (Cell Signaling) (2589); Anti-GFP (abcam) (ab1218); Anti-Presenillin-1 (Cell Signaling) (5643); Anti-Pecam1 (R&D Systems) (AF3628); Anti-Cd68 (Invitrogen) (14-0681-82); Cy3^TM^-conjugated Anti-SMA (Sigma) (C6198);

### Cell Culture and Treatments

Primary HUVECs were obtained from the Yale Vascular Biology and Therapeutics core facility and were used in all the *in vitro* experiments except Figure 3G. Each batch contains cells pooled from three donors. Primary HAECs were obtained from Lonza (Lonza: cc-2535). Cells were cultured in M199 (Gibco: 11150-059) supplemented with 20% FBS, 1x Penicillin-Streptomycin (Gibco: 15140-122), 60 µg/ml heparin (Sigma: H3393), and endothelial growth cell supplement (ECGS; the total formulation termed hereafter as complete medium). HUVECs were used for experiments between passages 3 and 6. For ligand-induced Notch activation, HUVECs were seeded onto the surface coated with 10µg/mL recombinant DLL4 (R&D: 1506D4050) as described^17^.

### Lentiviral transduction

Lenti-X 293T cells (Clontech, 632180) were cultured for at least 24 hours in DMEM supplemented with 10% FBS and lacking antibiotics, then transfected with lentiviral plasmids encoding the gene of interest and packaging plasmids (Addgene: 12259 and 12260) using Lipofectamine 2000 (Thermo Fisher Scientific: 11668-019) following the manufacturer’s protocols with Opti-MEM medium. Conditioned media from these cultures were collected 48 hours later, sterilized through 0.45µm filters and added to HUVECs together with 10µg/ml of polybrene (Sigma: 107689). After 24 hours, cells were switched to complete medium for 48 hours.

### siRNA transfection

HUVECs were cultured in EGM^TM^-2 Endothelial Cell Growth Medium-2 BulletKit^TM^ (Lonza: CC-3156 and CC-4176) for 24 hours before transfecting with RNAiMax (Thermo Fisher Scientific: 13778-150) with 20nM siRNA in Opti-MEM (Gibco: 31985-070) according to the manufacturer’s instructions. After 6 hours, cells were switched to EGM-2 medium and used for experiments 3 days later. Gα protein siRNAs were custom designed based on previous publications^44–46^. Latrophilin knockdown used ON-TARGET plus Smartpool siRNAs from Dharmacon (LPHN1: L-005650-00-0005; LPHN2: L-005651-00-0005; LPHN3: L-005652-00-0005). Similarly, PIEZO1, ARRB1 and ARRB2 knockdowns were achieved using Dharmacon SmartPools (PIEZO1: L2-020870-01-0005; ARRB1: L-011971-00-0005; ARRB2: L-007292-00-0005).

### Shear stress

HUVECs were seeded on tissue culture-treated plastic slides coated with 10 µg/ml fibronectin for overnight at 37°C and grown to confluence. For short-term experiments, cells were starved over-night in M199 medium with 2% FBS and 1:10 of ECGS. Slides were transferred to parallel flow chambers and shear stress applied as described^47^.

### Removal of membrane cholesterol by β-cyclodextrins

To deplete membrane cholesterol, cells were treated with methyl-β-cyclodextrin (MβCD) in serum-free medium as described before^48^ at the concentrations and for the durations indicated in the corresponding figures. For hydroxypropyl-β-cyclodextrin (HPβCD) treatment, cells were incubated in complete medium as described^36^.

### RNA isolation and RT-PCR

RNA was isolated from HUVECs using RNeasy kit according to the manufacturer’s instructions and quantified using a nanodrop spectrophotometer. Following cDNA synthesis using Bio-Rad iScript kit, RT-PCR was performed as follows: each PCR reaction contains 42 two-step amplification cycles consisting of: (a) denaturation at 95°C for 30s, and (b) annealing and extension at 60°C for 30s. The amplification curve was recorded and the relative transcript level of the target mRNA in each sample was calculated by normalization of Ct values to the reference RNA (GAPDH or 18S). Primer sequences are shown in Table 1.

**Table 1:**
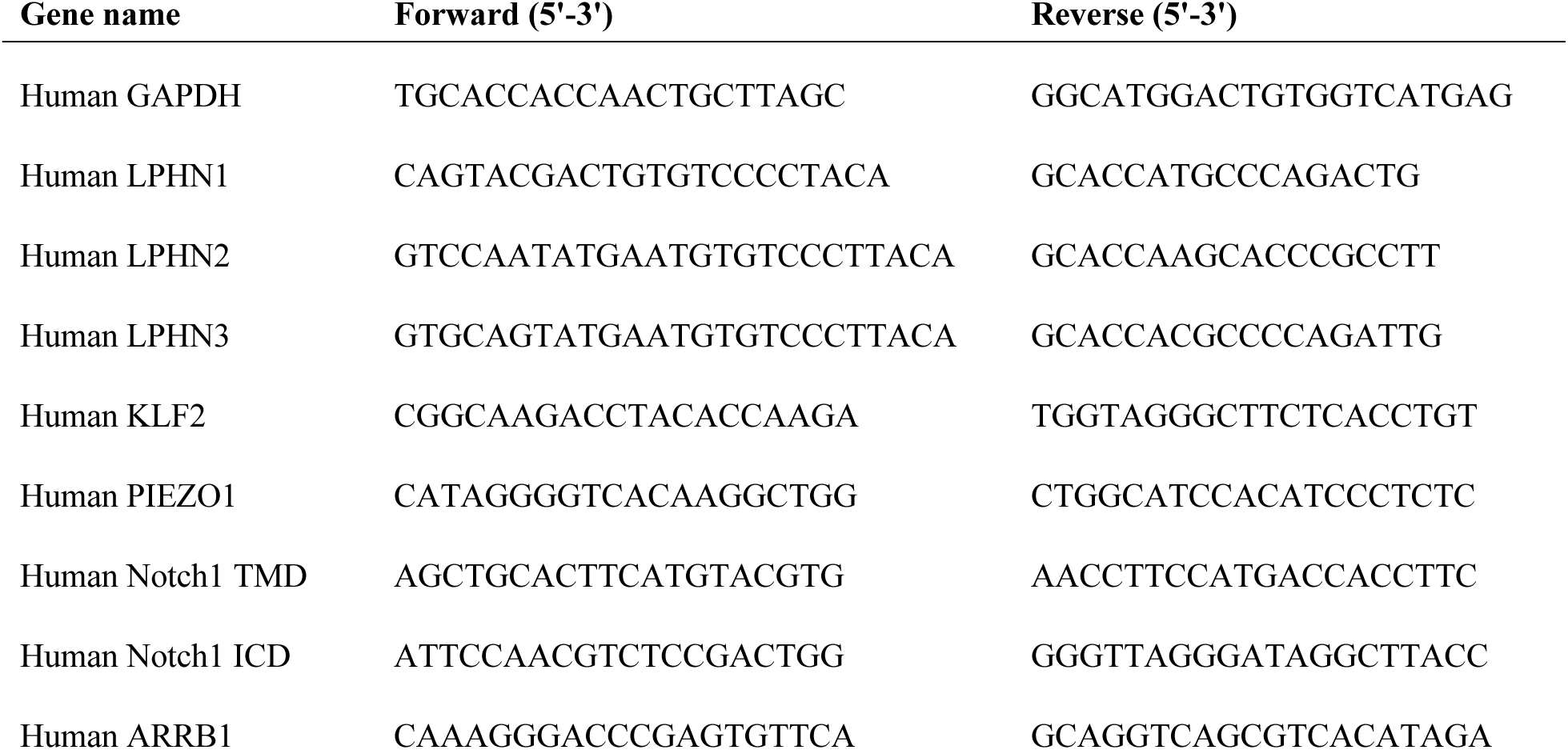
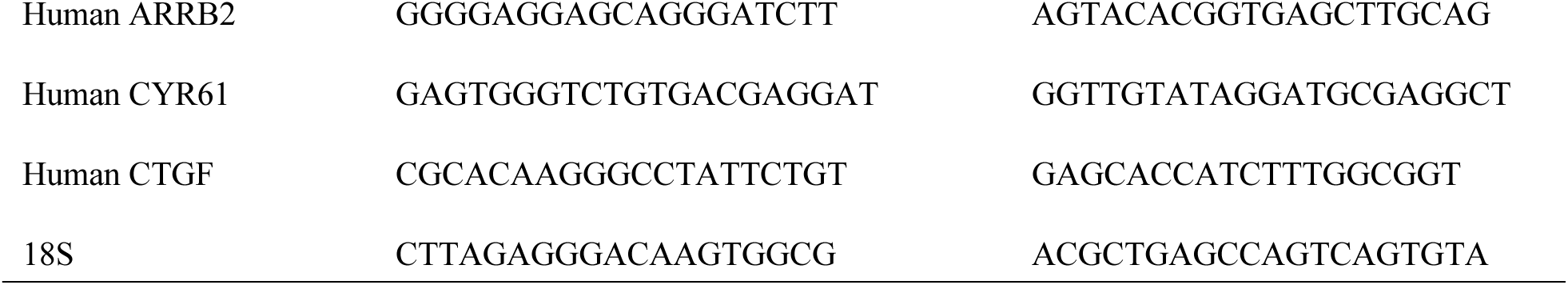
Primer sequences used for quantitative PCR in this study.

### RNA Sequencing

RNA was extracted from HUVECs subjected to Lphn2 knockdown and/or Notch1 domain over-expression or from HUVECs subjected to Piezo1 knockdown using QIAGEN RNEasy kit following the manufacturer’s protocol. RT-PCR confirmed efficient gene knockdowns and successful overexpression of Notch1 TMD (this study) or Notch1 ICD (addgene; 64622), and robust flow-mediated induction of KLF2, a previously validated Lphn2-independent flow-responsive gene^5^. Libraries were prepared based on poly A enrichment and sequenced using standard protocols of Yale center of Genomic Analysis.

### Data analysis of RNA Sequencing

Raw RNAseq data in FASTQ format were first assessed for quality using FastQC (v0.12.1). Adapter sequences and low-quality bases were trimmed from the reads using fastp (v0.23.2) with default parameters. The resulting high-quality reads were aligned to the human reference genome (GRCh38) using STAR aligner (v2.7.11a). PCR duplicates were identified and removed using Pi-card MarkDuplicates (v2.25.6) to ensure accurate quantification. Gene-level read counts were generated with the featureCounts function from the Subread package (v2.0.3) using GENCODE gene annotation.

To correct for potential batch effects, count matrices were processed with ComBat-seq in Bioconductor sva package (v.3.56.0). Count normalization was conducted using edgeR (v.4.6.2) and differential gene expression analysis was performed using DESeq2 (v1.44.0) with significance thresholds set at false discovery rate (FDR) < 0.01 and |log2 fold change| > 1. Gene Ontology (GO) and pathway enrichment analyses were conducted using the genekitr package (v1.0.5) to identify significantly enriched biological processes and pathways among differentially expressed genes.

A scatter plot comparing *LPHN2* KD vs. *PIEZO1* KD differentially expressed genes (scrambled_PSS vs. scrambled_static) was generated for genes passing an |log2FC| > 1 threshold on at least one axis, with points colored red/blue/purple for genes meeting the threshold on both axes and belonging to the *LPHN2*-dependent, *PIEZO1*-dependent, or overlapping gene sets (gray otherwise), and marker size scaled to -log_10_(adjusted p-value). A companion Venn diagram was created using matplotlib-venn to show the overlap between *LPHN2*-dependent and *PIEZO1*-dependent DEGs, with the same color annotation as in the scatter plot and a surrounding box annotated with the total gene count to represent genes outside both sets.

### Single-cell RNA sequencing analysis of public human atherosclerotic carotid artery data

Single-cell RNA sequencing data from human carotid artery control and atherosclerotic plaque samples were retrieved from Zenodo (https://zenodo.org/records/17520898). Analysis employed Scanpy (version 1.10.4), including generation of uniform manifold approximation and projection (UMAP) visualization for cell-type annotation and comparison of genes-of-interest expression between control and plaque tissues.

### Western Blotting

Cell lysates or immunoprecipitation samples extracted in Laemmli sample buffer were separated by SDS-PAGE and transferred onto nitrocellulose membranes. Membranes were blocked with 5% milk in TBS-T and probed with primary antibodies at 4°C for overnight. The targeting proteins were visualized using HRP-conjugated secondary antibodies and the HRP-luminol reaction.

### Immunofluorescence

HUVECs were washed with PBS, fixed for 10 minutes with 4% PFA in PBS and permeabilized with 0.5% Triton X-100 in PBS for 10 minutes, and blocked with 3% BSA in PBS for 1 hour. Slides were then incubated with primary antibodies at 4°C for overnight, washed with PBS and incubated with secondary antibodies and Hoechst for 1 hour. Samples were then washed 4 times with PBS and mounted.

### Image analysis for nuclear signal quantification

Images were acquired on a PerkinElmer spinning disk microscope using a 20x objective. Hes1 and YAP1 nuclear translocation was quantified using a custom ImageJ macro code. The code first creates nuclear masks derived from Hoechst-stained images, then generates cytoplasmic masks by enlarging these nucleus masks using ImageJ’s built-in ENLARGE function, performing this expansion five times and subsequently subtracting the original nucleus masks to isolate the cytoplasmic region. The code quantifies the intensity of Hes1 or YAP1 images within both the nucleus and the cytoplasm areas, reporting the nuclear intensity for Hes1/YAP1 or calculating the ratio of nuclear to cytoplasmic intensity of YAP1.

### Coimmunoprecipitation of Lphn2 for detection of interaction with Notch components

Cells were infected with lentivirus encoding Lphn2-mClover3, at 3 days after infection washed with cold PBS, scraped, and homogenized in 200 μl of buffer containing 10 mM Tris-HCl, pH 7.5, 5 mM MgCl2, 1 mM DTT, 0.25 M sucrose, and a cocktail of protease inhibitors (Thermo Fisher Scientific, 78440) per 1 well of 6 well plate. Nuclei and unbroken cells were removed by centrifugation at 500 × g for 5 min at 4°C, and the postnuclear supernatant was centrifuged at 100,000 × g for 1 hour at 4°C to separate the membrane and cytoplasmic fractions. Membrane fractions were solubilized overnight at 4°C in PIPES buffer with 1% CHAPSO (10 mM PIPES, pH 7.0, 5 mM CaCl_2_, 5mM MgCl_2_, 1% CHAPSO, and protease inhibitor cocktail). After solubilization and centrifugation to remove undissolved fraction, membrane fraction lysates were incubated with GFP-Trap^TM^ beads (Chromotek, gta) for 2 hours at 4°C with gentle rotation. Beads were washed three times with PIPES buffer containing 0.25% CHAPSO, eluted with 1x Laemmli buffer and analyzed by SDS-PAGE and immunoblotting targeting Presenillin-1 or Notch1 using the protocol described above.

### Proximity ligation assay

Proximity Ligation assay targeting GFP and Presenillin-1 was performed using the Duolink In Situ PLA kit from Sigma-Aldrich, following the manufacturer’s protocol as described in the Duolink Fluorescence User Manual (https://www.sigmaaldrich.com/US/en/technical-documents/proto-col/protein-biology/protein-and-nucleic-acid-interactions/duolink-fluorescence-user-manual). Briefly, cells were cultured on ibidi µ-Slide 15 Well 3D chambered coverslips (ibidi, cat no.: 81506) to enable staining in a small volume. Following fixation and permeabilization, cells were blocked with Duolink Blocking Solution and incubated overnight at 4°C with primary antibodies (GFP: 1:100, mouse, abcam, ab1218; Presenillin-1: 1:100, rabbit, Cell Signaling, 5643). Species-specific PLA probes (anti-mouse PLUS and anti-rabbit MINUS) conjugated to complementary oligonucleotides were then applied, followed by ligation and rolling-circle amplification using fluorescently labeled oligonucleotide probes, generating discrete fluorescent puncta at sites where the two proteins were in close proximity (<40 nm). Nuclei were counterstained with DAPI, and slides were kept in anti-FADE medium prior to imaging.

Images were acquired on a Leica SP8 confocal microscope using Leica Application Suite software. PLA signal quantification was performed as previously described^49^, using ImageJ/Fiji for puncta detection and normalization.

### Animals

All mice used in this study were on the C57BL/6J background. B6;129S6-Adgrl2^tm1Sud^/J mice (Jackson number 023401, hereafter Lphn2^flox/flox^) and B6;129-Tg (Cdh5-Cre)1Spe/J mice (Jackson number 017968, hereafter CDH5-Cre^ERT2^). Lphn2^flox/flox^ mice were crossed with CDH5-Cre^ERT2^ to get Lphn2^flox/flox^; CDH5-Cre^ERT2^ mice (hereafter Lphn2 iECKO). Lphn2^flox/flox^ mice (also treated with tamoxifen) served as controls. Mice were maintained in a light- and temperature-controlled environment with free access to food and water. Consistent with federal guidelines, all animal usage was approved by the Yale University Institutional Care and Use Committee.

### Immunostaining and imaging of mouse aorta

Aortae were perfusion-fixed in situ with 4% paraformaldehyde (overnight) prior to staining with primary Hes1 antibody (Cell signaling: 11988) or primary Yap antibody (Cell signaling: 14074) and fluorophore-conjugated secondary antibody (Invitrogen, donkey anti-Rabbit IgG (H+L) Highly Cross-Adsorbed Secondary Antibody, Alexa Fluor 647, A31573) in Claudio buffer (1% FBS, 3% BSA, 0.5% Triton X-100, 0.01% Sodium deoxycholate, 0.02% sodium azide in PBS pH7.4)^50^. Cells were stained with Hoechst (Invitrogen: H1399) to label nuclei. Images of stained vessels were acquired using a Leica SP8 confocal microscope with the Leica Application Suite (LAS) software.

### DAPT injections

DAPT dissolved in 5% DMSO, 40% PEG300, 5% Tween 80, and 50% deionized water was injected intraperitoneally into 8-week-old mice at a dose of 50 mg/kg daily for 5 consecutive days.

### Mouse atherosclerosis model

Mice at 8-9 weeks of age were injected intraperitoneally with murine Pcsk9 adeno-associated virus (pAAV/D377Y-mPcsk9; 10^11^ copies, produced by the Gene Therapy Program Vector Core at the University of Pennsylvania School of Medicine or Vector Biolabs). Mice were then fed a high fat diet (Clinton/Cybulsky high fat rodent diet with regular casein and 1.25% added cholesterol, D12108c, Research Diet) for 8 weeks before sacrifice.

### Mouse whole aorta *en face* oil red O staining

Mice were euthanized with an overdose of isoflurane (Henry Schein) and perfused through the left ventricle with PBS followed by 3.7% formaldehyde. The whole aorta was carefully dissected and fixed in 3.7% formaldehyde overnight at 4℃, then washed 3x with PBS. For whole aorta Oil Red O staining, the formaldehyde fixed whole aorta was opened longitudinally, pinned *en face* on a soft bottom dish, briefly rinsed with 78% methanol, then incubated with Oil Red O solution (0.2% Oil Red O in 7:2 methanol:1M NaOH). Stained tissue was washed 3x in 78% methanol and photographed using a Leica DFC295 digital microscopic camera.

### Mouse blood collection and measurement

Blood samples were collected from mice by heart puncture at sacrifice. Blood was mixed with EDTA to prevent coagulation and was then centrifuged at 14000rpm at 4℃ for 10 minutes and the supernatant serum removed carefully by pipet. Total serum cholesterol was determined using a cholesterol oxidase/colorimetric assay (Abcam; #65390). Plasma TAG levels were measured using a commercially available kit (Wako Pure Chemicals).

### Tissue staining of the aortic root

Aortic root sections were dissected and fixed in 4% formaldehyde overnight at 4°C. Tissues were then washed with PBS three times and cryoprotected in 30% sucrose in PBS for 24 h, embedded in optimal cutting temperature (OCT) compound, and frozen on dry ice. Tissue blocks were cut into 10μm sections using a cryostat (Leica) and sections were stored at −80°C until use.

Hematoxylin and eosin (H&E) staining and Masson’s trichrome staining on OCT sections were performed by Yale’s Research Histology Laboratory using standard protocols, and images were acquired using a Nikon 80i microscope.

For immunofluorescence, OCT sections were thawed for 15 minutes, outlined with a hydrophobic pen and air-dried for additional 15 minutes. Sections were then permeabilized and blocked with 3% FBS with 0.3% Triton X-100 in PBS (prepared with nuclease-free water; hereafter, perm/block buffer) for 1 h at room temperature, followed by overnight incubation at 4°C with primary anti-bodies (anti-Cd68, 1:200, Invitrogen, 14-0681-82; anti-Pecam1, 1:100, R&D Systems, AF3628) in perm/block buffer. Sections were then washed three times with perm/block buffer and incubated with secondary antibodies diluted 1:1000 in perm/block buffer, along with Hoechst 33342 (1:1000, Invitrogen, H1399) and Cy3^TM^-conjugated alpha-SMA (1:100, Sigma, C6198) for 1 h at room temperature. Secondary antibodies (Invitrogen) were donkey anti-rat, or -goat conjugated with Alexa Fluor 488, or 647, respectively. Slides were washed three times with perm/block buffer and mounted in DAPI Fluoromount G (Southern Biotech; 0100–20). Images were acquired on a Leica SP8 confocal microscope using Leica Application Suite software.

### Masson’s Trichrome staining analysis

Masson’s Trichrome–stained sections were analyzed using color deconvolution function built in ImageJ/Fiji. A custom color vector (Color 3: R = 0.59227383, G = 0.3264422, B = 0.7366459) was applied to isolate the collagen-specific channel. Within a defined region of interest in the aortic root, the mean grayscale intensity of the collagen channel was measured and used as an index of collagen content.

### Statistical Analysis

Data are presented as means ± SEM. Statistical significance was determined using Student’s t-test or ANOVA as indicated in the figure legends. P < 0.05 was considered significant.

## Acknowledgements

This work was supported by NIH grant R01 HL155543 and RO1 HL169510 to MAS.

## Author Contributions

KT and MAS designed the project. KT performed *in vitro* experiments, *in vivo* mouse experiments and bioinformatics analysis. MChen performed *in vivo* mouse experiments. DJ helped with mouse dissections. MChanduri helped with *in vitro* experiments. YMK performed *in vitro* experiments and provided technical support for *in vivo* and *in vitro* experiments. XW assisted with mouse dissections and tissue sectioning. HZ performed measurements of blood lipid levels from atherosclerosis mouse experiments. XZ carried out some flow experiments. JS performed tissue staining and imaging. CFH provided equipment and advice for atherosclerosis work. KT wrote the first draft of the manuscript. KT and MAS wrote and reviewed the manuscript with the input of all authors.

---

**Supplemental Figure 1:**
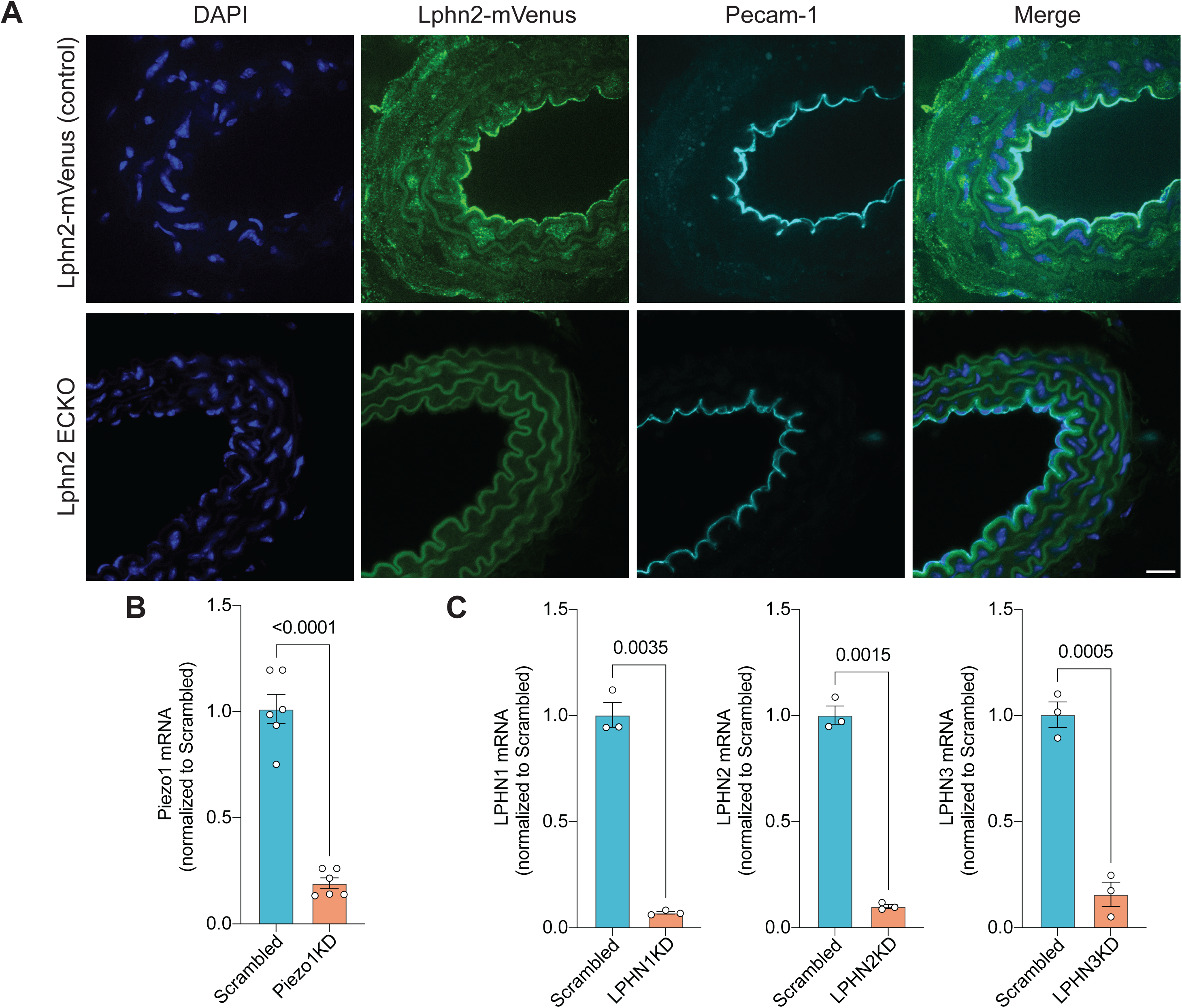
Confirmation of Lphn2 endothelial-cell knockout (ECKO) and knockdowns used in this study. (**A**) Representative images of carotid artery sections stained for DAPI and Pecam1 (endothelial marker), along with fluorescence of Lphn2-mVenus expressed from the endogenous Lphn2 promoter. Control arteries showed robust endothelial Lphn2-mVenus expression, which was absent in after Lphn2 ECKO. Scale bar: 20µm. (**B**) Quantitative PCR for Piezo1 mRNA in Scrambled and Piezo1 KD in HUVEC. Values are normalized to Scrambled. Statistics used unpaired Student’s t-test. (**C**) Quantitative PCR for Lphn1 (left), Lphn2 (middle) and Lphn3 (right) mRNA. Values are normalized to Scrambled. Statistics used unpaired Student’s t-test.

**Supplemental Figure 2:**
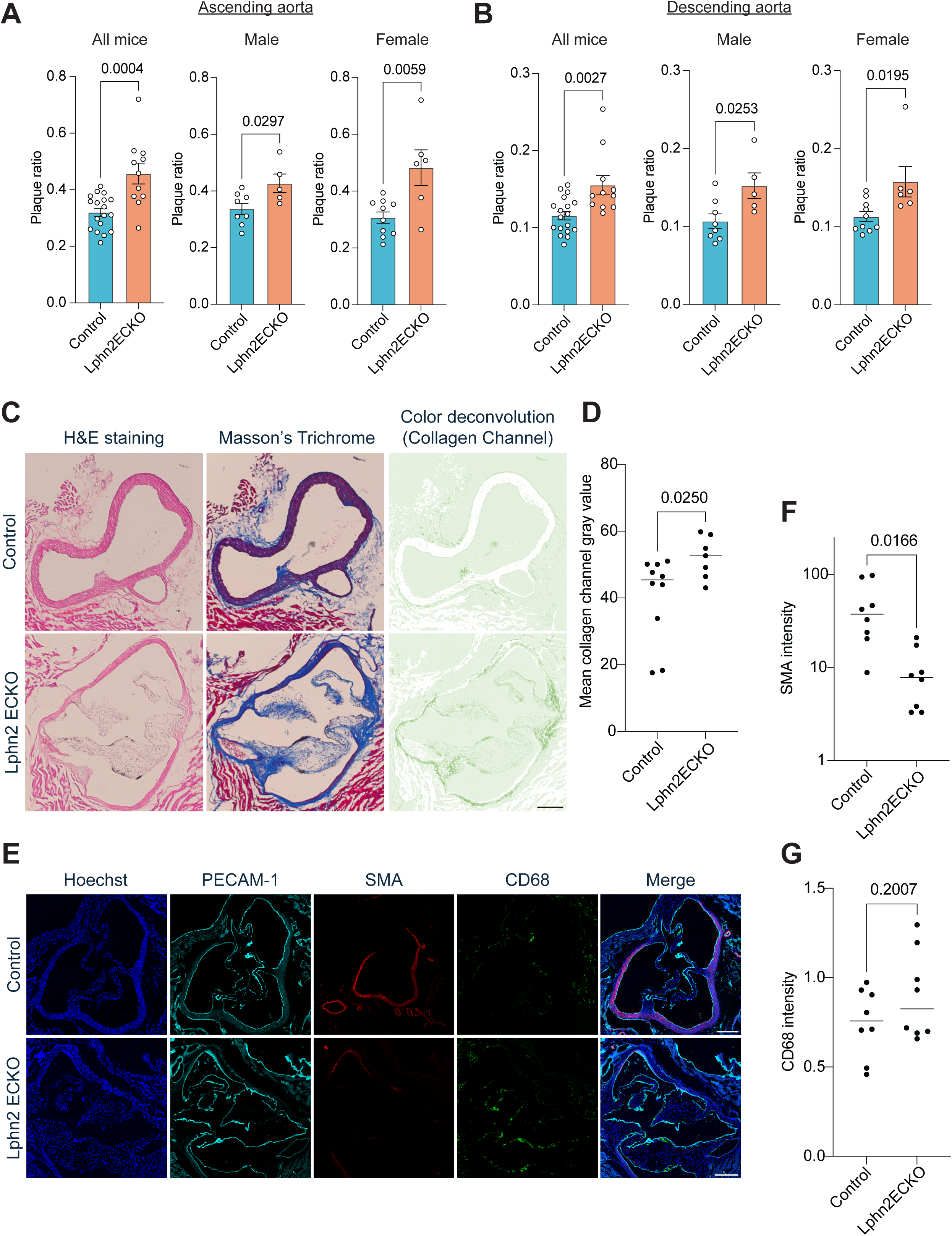
Characterization of atherosclerotic plaques of Lphn2 ECKO mice: (**A**) Quantification of atherosclerotic plaque area in the ascending aorta in Lphn2 ECKO mice versus control mice (ratio plaque area to total area). Data are from the same datasets as Figure 1C. Statistics were calculated using unpaired Student’s t-test. (**B**) Quantification of atherosclerotic plaque area in the descending aorta, as in (**A**). (**C**) hematoxylin and eosin (H&E) staining, Masson’s trichrome staining and color deconvolution of Masson’s trichrome images of aortic root sections from control and Lphn2 ECKO mice. Scale bar: 200µm. (**D**) Quantification of collagen signal intensity in aortic root regions in control and Lphn2 ECKO mice. Statistics used unpaired Student’s t-test. (**E**) Representative images of aortic root sections stained for DAPI, Pecam-1, SMA and Cd68. Scale bar: 200µm. (**F**) Quantification of SMA signal intensity in the aortic root. Statistics used unpaired Student’s t-test. (G) Quantification of Cd68 signal intensity in the aortic root. Statistics used unpaired Student’s t-test.

**Supplementary Figure 3.**
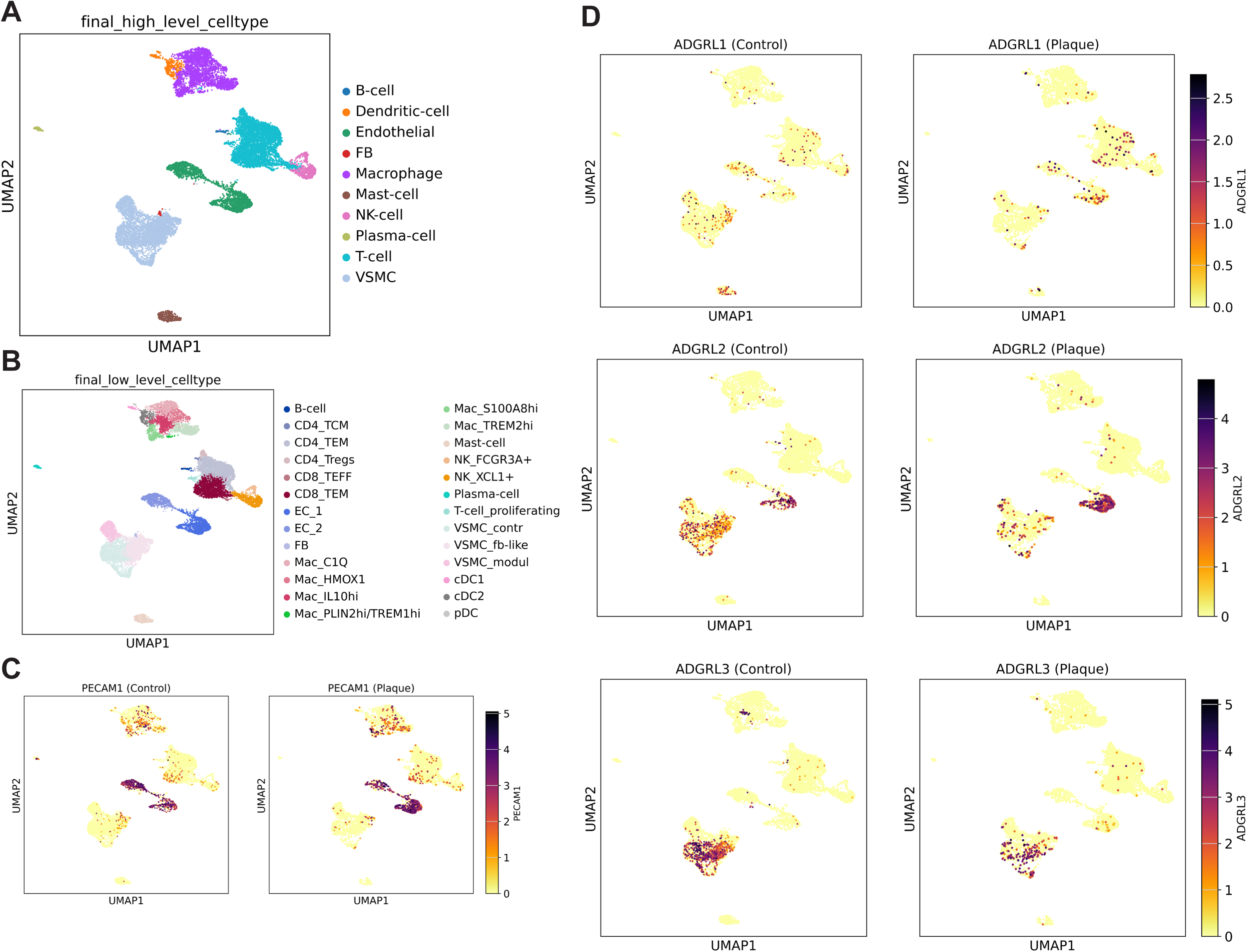
Expression of latrophilins in public single-cell RNA-sequencing data from human atherosclerotic plaques. (**A**) Low-resolution cell-type annotation. (**B**) High-resolution cell-type annotation. (**C**) Expression of PECAM1, an endothelial-cell marker. (**D**) Expression of LPHN1 (latrophilin-1). (**E**) Expression of LPHN2 (latrophilin-2). (**F**) Expression of LPHN3 (latrophilin-3).

**Supplemental Figure 4:**
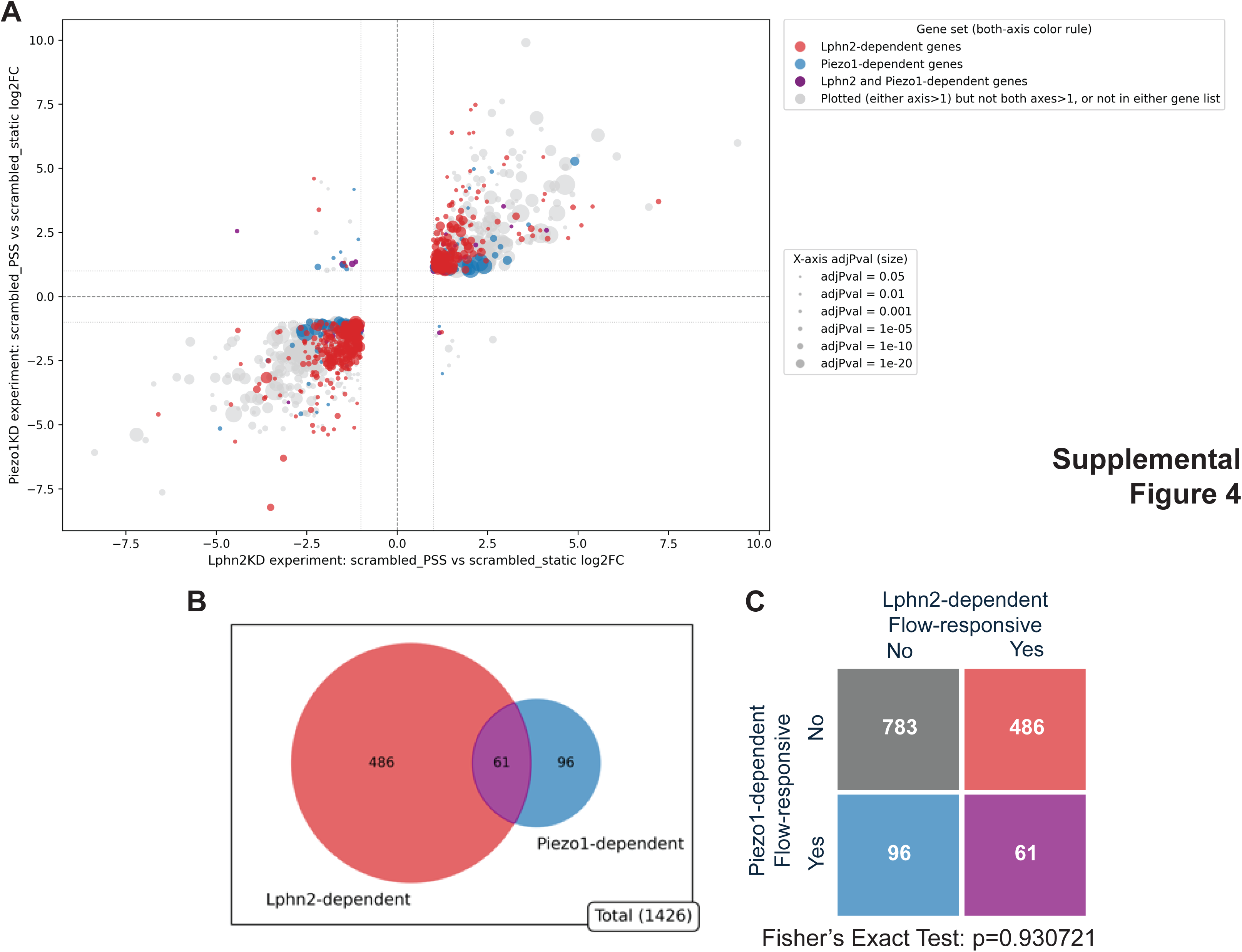
Flow-dependent genes that require Piezo1 vs Latrophilin-2. (**A**) Scatter plots of LPHN2-dependent and PIEZO1-dependent flow-responsive differentially expressed genes (DEGs). LPHN2-dependent DEGs are shown in red, PIEZO1-dependent DEGs in blue, and genes dependent on both are shown in purple. Dot size reflects adjusted p-value. (**B**) Venn diagram of DEGs between LPHN2 knockdown and PIEZO1 knockdown. (**C**) 2×2 table of DEGs between LPHN2 knockdown and PIEZO1 knockdown. Fisher’s exact test shows no significant association between LPHN2-dependent and PIEZO1-dependent gene profiles.

**Supplemental Figure 5:**
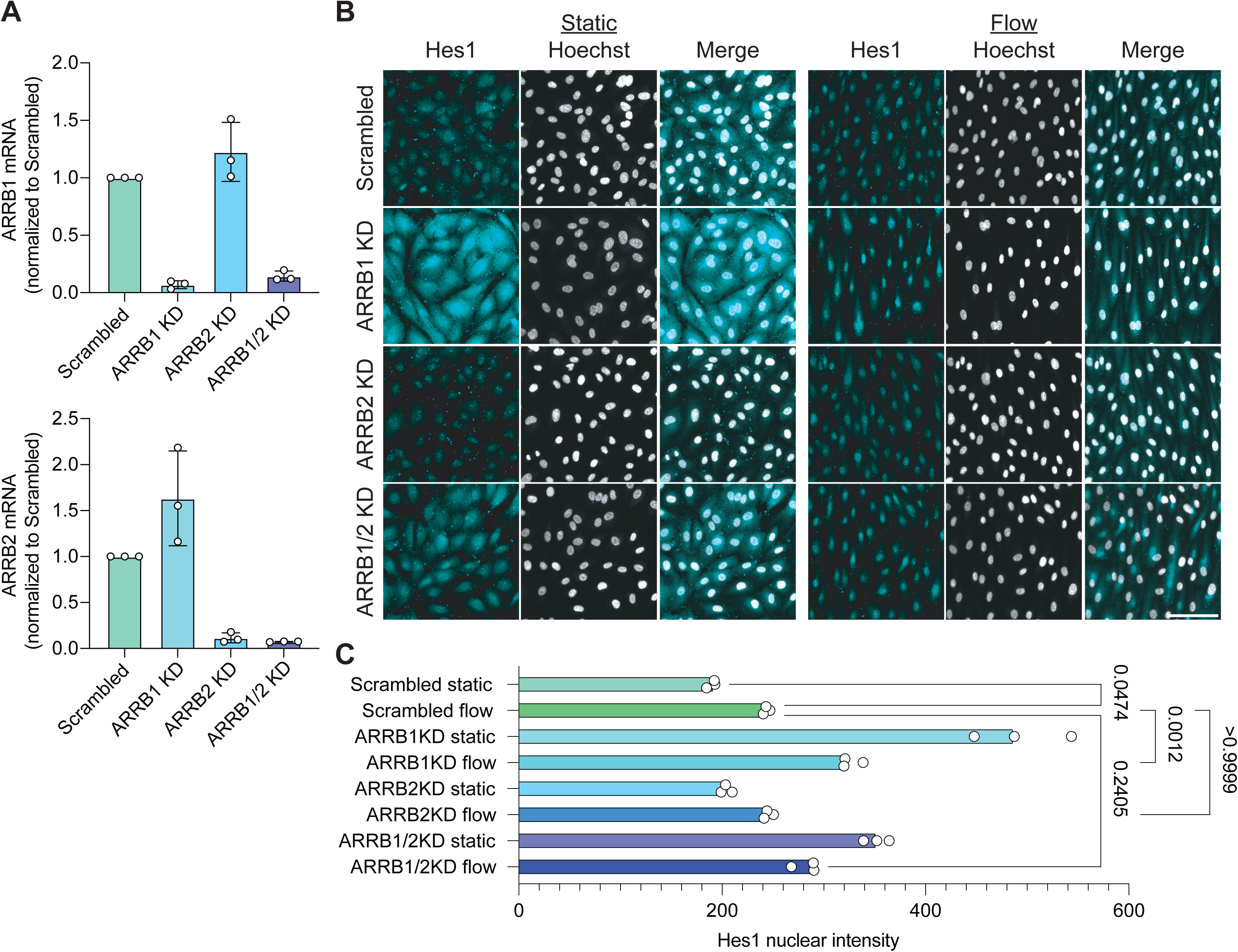
Beta-arrestins do not regulate flow-induced Notch signaling. (**A**) Quantitative PCR for ARRB1 (top) and ARRB2 (bottom) mRNA in Scrambled, ARRB1 KD, ARRB2 KD and ARRB1/2 KD in HUVECs. Values are fold change relative to Scrambled control. (**B**) Representative images of Hes1 and Hoechst staining in HUVECs following knockdown of beta-arrestin isoforms with or without flow. Scale bar: 50µm. (**C**) Quantification of Hes1 nuclear intensity from (**B**). Quantifications are from three independent experiments. Statistics using one-way ANOVA with Tukey post-hoc tests.

**Supplemental Figure 6:**
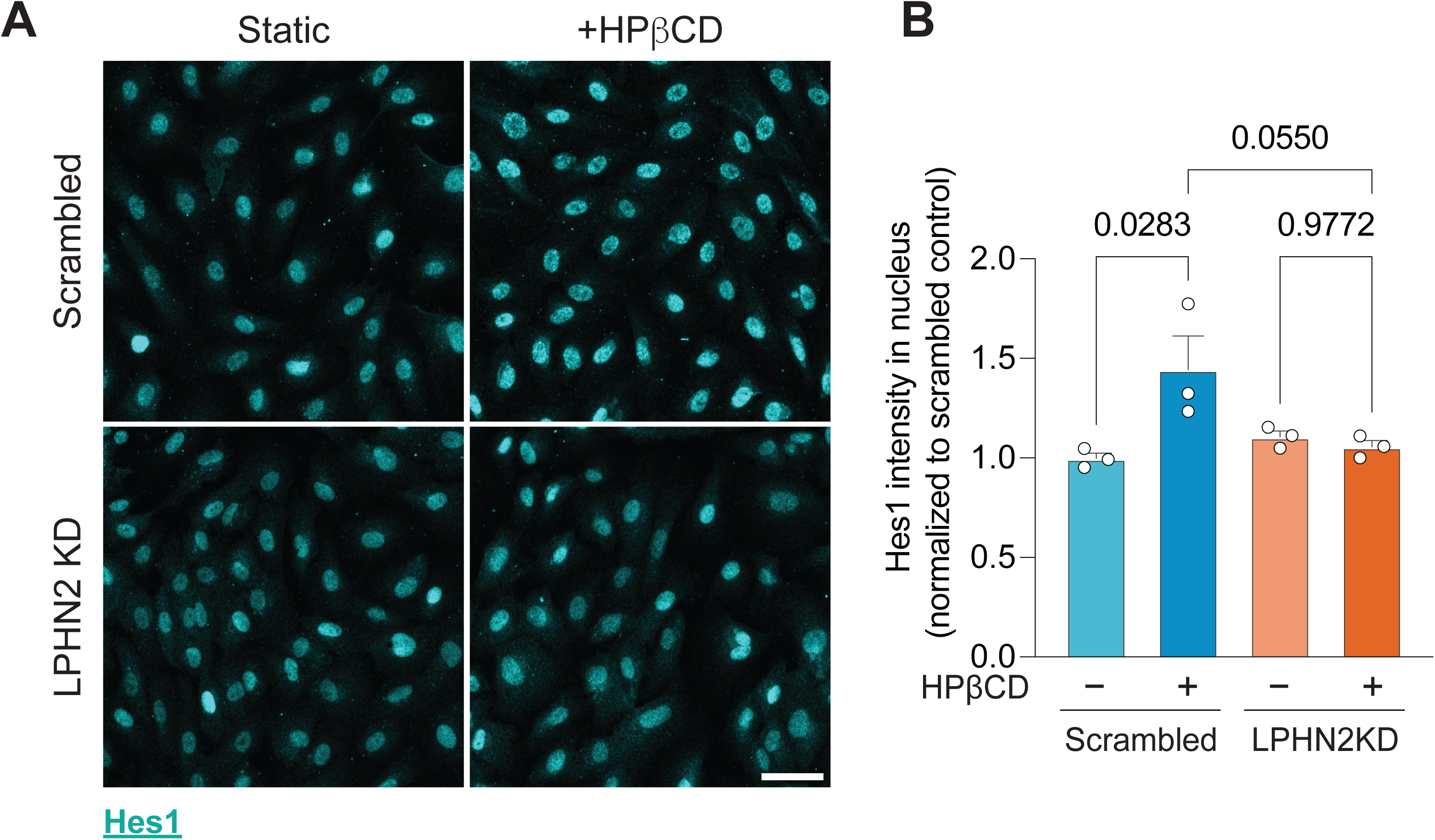
Removing membrane cholesterol induces the expression of Notch target gene Hes1. (**A**) Representative images of Hes1 with/without 5mM hydroxopropyl β-cyclodextrin for 2 hours. Scale bar: 50µm. (**B**) Quantification of nuclear Hes1 intensity from (**A**). Data are from three independent experiments. Statistics used one-way ANOVA with post-hoc Tukey test.

**Supplemental Figure 7:**
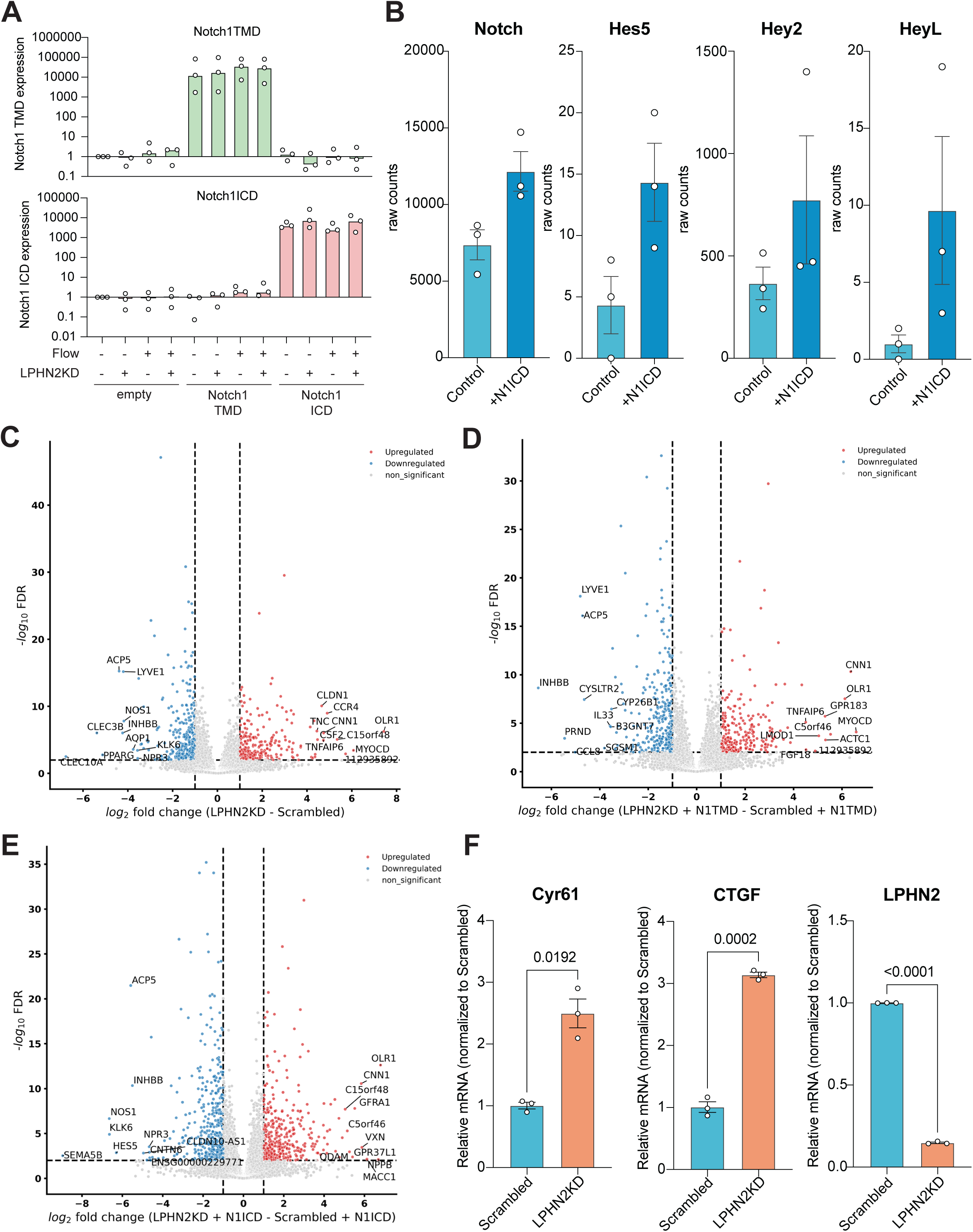
Validation of RNAseq experiments with rescue of LPHN2 knock-down by Notch1 constructs. (**A**) Quantitative RT-PCR of Notch1 TMD and Notch1 ICD in the samples analyzed in Figure 6. (**B**) RNA-seq read counts for Notch1 and Notch signaling target genes following N1ICD overexpression. (**C**) Volcano plot for differentially expressed genes between LPHN2KD and control siRNA. (**D**) Volcano plot for differentially expressed genes between LPHN2KD and control siRNA with overexpression of Notch1 TMD. (**E**) Volcano plot for differentially expressed genes between LPHN2KD and control siRNA with overexpression of Notch1 ICD. (**F**) Quantitative RT-PCR of canonical YAP target genes and Latrophilin-2 under Latrophilin-2 knockdown. Statistics were calculated by unpaired Student’s t-test.

**Supplemental Figure 8:**
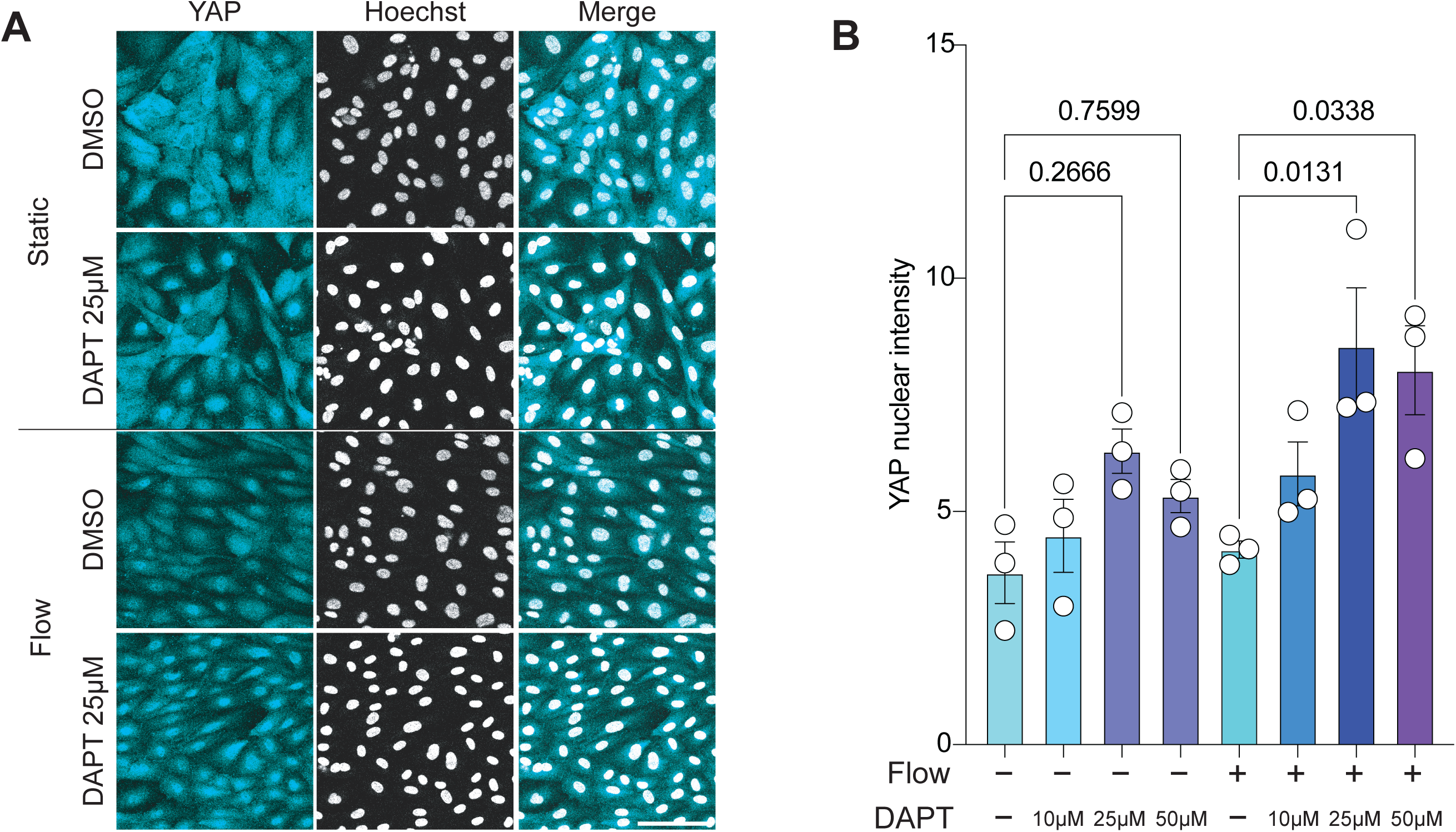
LPHN2-Notch axis induces vascular inflammation through YAP. (**A**) Representative immunofluorescence image showing increased YAP nuclear translocation ± 12 dynes/cm2 flow for 16 hours, with DAPT at the indicated doses. Scale bar: 50µm. (**B**) Quantification of YAP nuclear intensity from (A). Data are from three independent experiments. Statistics used one-way ANOVA with Tukey post-hoc tests.

**Supplemental Figure 9:**
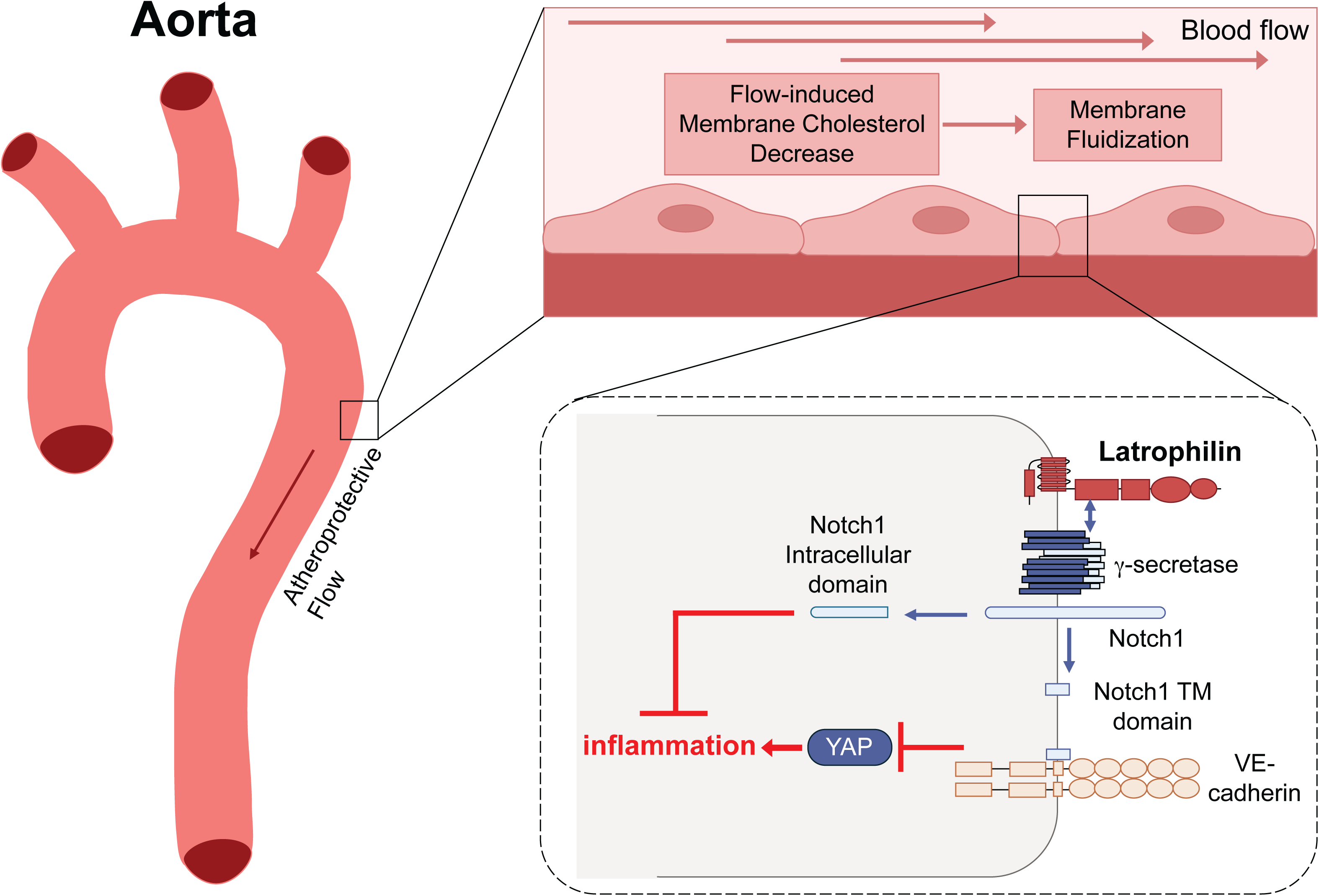
Pathway diagram for the Latrophilin-Notch axis Lphn2 mediates Notch1 activation in regions of physiological laminar shear stress. Both the N1ICD and the N1TMD contribute to suppression of inflammatory pathways, such as Yap, that promote atherosclerosis.

